# Proteome-wide multi-omics profiling of osteosarcoma transcription factor networks

**DOI:** 10.64898/2026.03.29.714917

**Authors:** Nguyen Xuan Thang, Emily Louise Beecher Martinsen, Mohamed Abdelhalim, The Trung Tran, Marit Ledsaak, Marie Rogne, Bernd Thiede, Ragnhild Eskeland

**Author notes:** Co-first authors (Nguyen Xuan Thang, Emily Louise Beecher Martinsen). Senior author (Ragnhild Eskeland).

## Abstract

Osteosarcoma (OS) is an aggressive bone cancer that most commonly affects children and young adults. OS exhibits a high degree of genomic complexity, as well as cellular plasticity, and dynamic transcriptional regulation is suggested to contribute to treatment resistance and metastasis. Cell lines are well characterized as models to advance our knowledge on OS biology. HOS and U2OS cells have increased invasiveness and higher migratory ability compared with MG63. In this study, we employed a tandem array of consensus transcription factor response elements (catTFREs) proteomic approach to characterize transcription factor (TF) regulatory networks related to OS aggressiveness. We mapped 7,594 proteins and enriched 352 transcription factors and coregulators. When we integrated proteomics with cell line specific gene expression and chromatin accessibility we classified the proteins into different OS cell line dependent sub-clusters and identified TFs and coregulators common for all cell lines and specific for individual cell lines. We demonstrate that RUNX2, MYBL2 and HMGA2 are specifically enriched in HOS and U2OS and may be linked to the cell aggressiveness. ETV5, JUNB, NFIX and ZEB1 were among TFs specific to MG63. Our analysis provides a more comprehensive understanding of the transcriptional drivers that shape OS regulatory landscapes and may have future therapeutic implications.

## INTRODUCTION

Osteosarcoma (OS) is a primary malignant mesenchymal tumour originating from bone or osteoid that accounts for approximately 20% of bone cancers [1]. There is a higher incidence of OS in children and adolescents, and the tumours tend to grow quickly with a high rate of metastasis. OS generally occurs in long bones and can be classified as low-, intermediate-, or high-grade malignancy based on biopsy and histology [2]. Standard treatment is chemotherapy and surgical resection and has not improved much over the past decade, however, one-third of OS patients do not respond to chemotherapy [3,4]. Due to OS rarity, genetic heterogeneity, and high genome instability, the aetiology of these tumours remains poorly understood. In terms of molecular genetics, the mutations or lack of activation of tumour suppressor genes, such as *Tumour protein 53 (TP53), Retinoblastoma 1 (RB1)*, and *Recq like helicase 4 (RECQL4)*, are key drivers of OS [5,6]. In addition, alterations in epigenetic factors *KMT2C (MLL3)*, *Alpha-Thalassemia/Mental Retardation Syndrome X-linked* (*ATRX)*, or oncogenic amplifications of *Runt*-*related transcription factor 2* (*RUNX2), Mouse double minute 2 (MDM2), Vascular Endothelial Growth Factor A (VEGFA), MYC*, and *Cyclin dependent kinase 4* (*CDK4)* are all drivers of OS [7–11]. Going beyond genomic alterations, transcription factors (TFs) may also contribute to OS heterogeneity by regulating cancer genomes. Here, studies of OS-derived cell lines that are highly proliferative and may represent malignant cancer characteristics can shed new light on tumorigenesis drivers [3].

TFs regulate target gene expression via binding to specific cis-regulatory DNA elements enriched in promoters and enhancers. TFs interact with co-activators or co-repressors to form regulatory complexes that orchestrate gene expression programmes [12]. Master TFs modulate chromatin shape and gene expression profiles to regulate cell fate [13]. While the TFs drive tissue- and cell-type identities [14,15], deregulation of specific TFs can drive cancer progression and metastasis [16–18]. Moreover, TFs account for over 20% of all oncogenes and were previously considered undruggable targets [19,20]. Deregulation of RUNX2 promotes metastasis and can function as a prognostic marker in OS, breast and colon cancer [21–23]. Furthermore, MYC acts as an oncogene in several types of cancers and is upregulated or amplified in many bone tumours, which is associated with a poor prognosis [24–26]. Mutations in TP53 are common and known oncogenic drivers in OS [27]. Overall, OS are complex tumours that have a strong tendency to metastasize and gain drug resistance, and genomic and transcriptomic analyses have had limited success in identifying interventions. A recent study by Sweet-Cordero and colleagues has defined two OS tumour epigenetic states by chromatin accessibility analysis [28]. The early osteoblast-derived state (EOD) is characterized by TFs RUNX2, MEOX2 and the MEF2 TF family and the late osteoblast-derived state (LOD) by FOSL and JUN TF families [28]. Based on single cell analysis, these two transcription states were shown to coexist in OS tumours and have different responses to drug treatment [28]. Thus, by generating maps of TFs and their DNA-binding activity in OS cells we may gain a better understanding of the underlying gene regulation and discover new druggable targets.

The generally low expression levels of TFs can pose challenges for detection and investigation, particularly at the protein level. In pursuit of overcoming this bottleneck, a pulldown approach with a DNA containing a concatenated tandem array of the consensus TFREs (catTFRE) has been developed to enrich TFs through their preferred interaction with DNA [15,29,30]. In this study, we employ a catTFRE approach in three OS cell lines (MG63, HOS and U2OS), identify 352 TFs and cofactors, and explore the regulatory networks and cellular characteristics for each cell line. By integrating the proteome mapped TF profiles with gene expression and chromatin accessibility, we identify a subset of TFs and cofactors that may be involved in orchestrating OS development and metastasis. Our catTFRE enriched factors are a resource that can be used to better understand transcriptional regulation OS metastasis and treatment resistance.

## RESULTS

### catTFREs enriches transcription factors independent of binding sites

TFs recognize and bind in specific DNA motifs with high affinity and selectivity. The catTFREs method integrates multiple canonical TF motifs based on TFs intrinsic sequence preferences to enhance the capture efficiency. Applications utilizing catTFREs have previously been shown to significantly improve the identification of TFs in cell lines and mouse tissues [15,29,30]. We developed a catTFRE approach [29] to enhance the detection of TFs in Osteosarcoma (OS) cell lines by MS analysis. OS cell lines have been classified into different subgroups based on their tumorigenic capacity (Table 1) [31]. We selected three OS cell lines for TF pulldowns that were less (MG63) and more aggressive (U2OS and HOS) based on their invasion and migration abilities [31,32]. For large scale production of linearized biotinylated DNA for pulldowns, we designed a three kilobase pair DNA array of 110 TFBS in duplicate (C1) from the JASPAR motif database [33] inserted into a vector (Figure 1A and Table S1A). C1 was compared to a control DNA fragment of similar size digested from the pGL4.26 vector (C4) which has a reduced number of TFBS (Figures 1 and S1A). When we applied CiiDER analysis [34] on C1 and control C4, we detected a total of 632 and 396 TFBS, respectively, with an overlap of 59% (Figures S1B and Table S1B). Both C1 and C4 were linearized by restriction digests, biotinylated on one end by Klenow and separated from the vector backbone by immobilization to M-280 Streptavidin Dynabeads^TM^ (Figure 1A, top). C1 and C4 bound to Dynabeads^TM^ were mixed with nuclear extracts (NE) from OS cell lines MG63, U2OS and HOS to enrich DNA binding proteins (Figure 1A, bottom, and Table S1C, four biological replicates). To compare the enrichment by catTFRE approach to total proteins in NEs we used liquid chromatography-MS (LC-MS) (Figures 1B, S1A,C,D and Table S1D). Pearson correlation analysis revealed a high correlation in protein abundance between cell lines, C1, C4, NE, and replicates (Pearson correlation coefficient R > 0.95) (Figure 1C). The protein counts in NEs were separated well into distinct groups compared with the catTFRE constructs C1 and C4, and also between cell lines (Figure 1C). In total, we identified 7,594 proteins across all replicates, with the highest number of proteins detected in the NEs from all three cell lines (Figure 1B). Nearly one-third (2,476 proteins) of the total proteins overlapped between all three cell lines and conditions (C1, C4 and NE; Figure S1F). When we filtered against a list of selected TFs and coregulators (Table S1E) [35], and mapped a total of 352 TFs across all three cell lines, find that the TF enrichment numbers varied for C1, C4, NE and replicates (Figure 1B and S1G, Table S1F). catTFREs C1 had a higher number of proteins compared to C4 with more than 175 TFs in all three replicates, however the overlap with TFBS in C1 and C4 was only 19 % and 11 %, respectively (Figures S1C and D). More than half of these (90 TFs) were identified in all three cell lines and conditions analysed by MS (Figure S1F). Using the Weighted Gene Co-expression Network Analysis (WGCNA) tool (Langfelder and Horvath, 2008), we analysed the protein data to cluster proteins identified in NE and enriched by C1/C4 in HOS, MG63 and U2OS. The heat map revealed a correlation of 10 clusters comprising a total of 7,594 proteins, detected across all conditions (Figure 1D and Table S1G). Most proteins were highly associated with the NE condition (4584 proteins), including the clusters II (1694 proteins), III (757 proteins), IV (611 proteins), V (803 proteins), and VI (719 proteins). Cluster IX had 291 proteins that were enriched by C1 and C4 catTFREs from all three OS cell lines. Interestingly, we mapped clusters specific to C1 and C4 catTFREs for each OS cell line, and this was clusters I (748 proteins) and VIII (292 proteins) from MG63, cluster VII (779 proteins) from U2OS, and cluster X (900 proteins) from HOS. RUNX2, Paired-like homeodomain transcription factor 1 (PITX1), and forkhead box O1 (FOXO1) are transcription factors shown to regulate bone development and progression of osteosarcoma [36–39]. According to MS counts, both C1 and C4 have enriched these TFs compared to the detection in NE, where for example RUNX2 was more highly enriched in HOS, paired-like homeodomain TF 1 (PITX1) was higher in MG63, and FOXO1 more prevalent in U2OS (Figure S1E).

**Figure 1.**
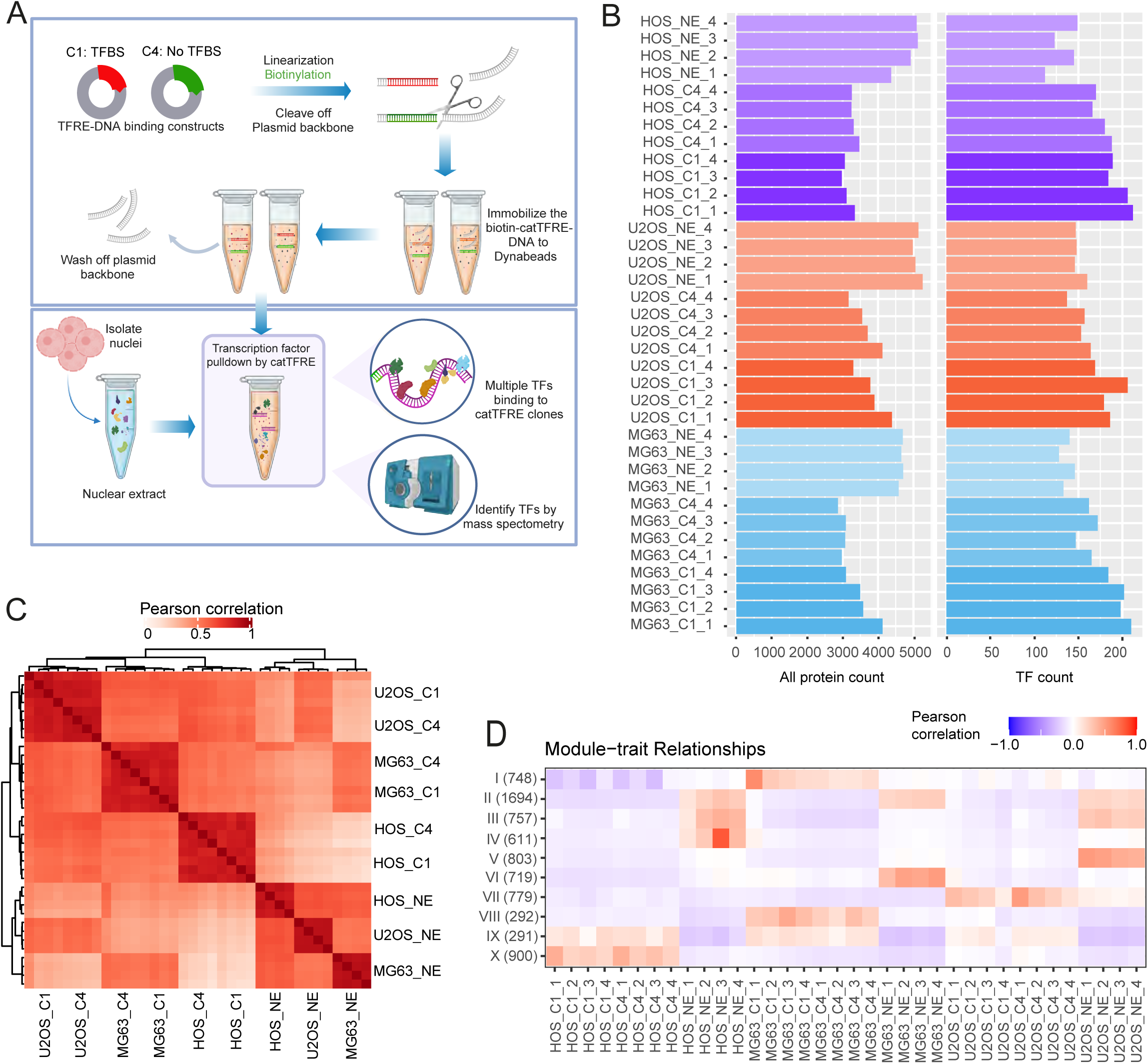
A catTFREs approach based on restriction digest and biotinylation of 3-kilobase insert sequences C1 and control C4 to enrich transcription factors from nuclear extracts. A) Schema of the concatenated tandem array of the consensus transcription factor response elements (catTFREs) workflow utilising two plasmids inserts C1 (designed with approximately 110 TF binding sites (TFBS) in duplicates with linkers) and C4 (reduced TFBS). Detailed TFBS are provided in Table S1A and B. Inserts were linearized by restriction enzyme digestion and biotinylated by Klenow enzyme. Biotinylated C1 and C4 were immobilized on streptavidin coated magnetic beads and incubated with nuclear extracts (NE) for detection of DNA binding proteins by mass spectrometry (MS). B) MS spectral counts of proteins enriched from NEs or by TFRE C1 and C4 pulldown. Four replicates with total count of proteins (left) and TFs only (right) from HOS (purple), U2OS (red) and MG63 (blue). All source data are provided in Table S1C-F. C) Heatmap representing the Pearson correlation of spectral counts by samples from HOS, U2OS and MG63. Scale represents low correlation in white to self-correlations as red-brown. D) Clustering of proteins by Weighted Gene Co-expression Network Analysis from MS spectral counts of NE and C1/C4 in HOS, MG63 and U2OS. Clusters I to X with the number of proteins represented by Pearson correlation across all conditions. Proteins are listed in Table S1G.

**Table 1.**
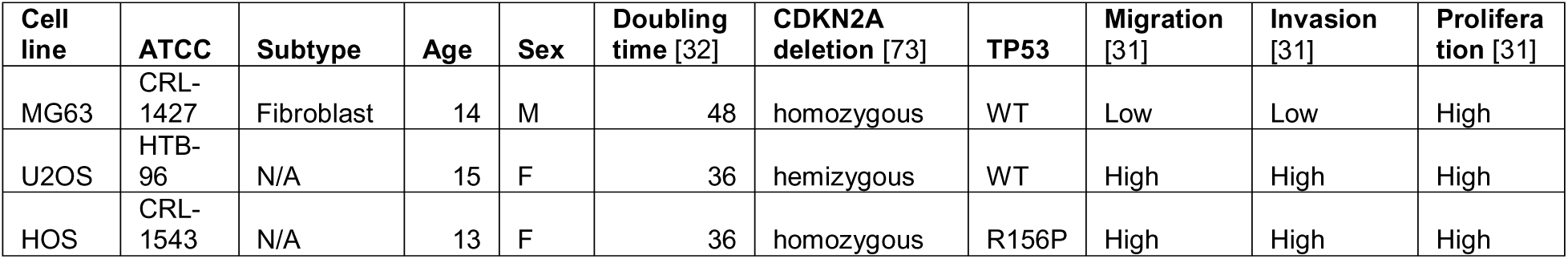
Overview of Osteosarcoma Cell lines.

We have shown that our catTFREs approach enriches TFs by MS from three different OS cell lines, which is in line with previous studies [29,40]. Moreover, we find that TFs bind linearized DNA independent of DNA sequence provided, but that there is some sequence preference as we observed a higher TF enrichment by C1 over control C4.

### catTFREs pulldowns enrich transcription factors from OS nuclear extracts

Next, we explored the differences between proteins detected in NEs versus the catTFREs approach in each cell line. Principal component analysis (PCA) of the proteins and mapped TFs by MS showed that the samples were correlated according to enrichment in NEs, catTFREs (C1 and C4), and OS cell lines (Figure S2A). We applied a stringent filtering to the MS data, selecting proteins that were detected in all replicates of at least one condition. Nonsupervised hierarchical clustering analysis revealed six distinct protein clusters (1-6) across all samples (Figure 2A). Three clusters represented proteins specific to OS cell lines, with 223 proteins representing U2OS (cluster 2), 196 proteins were more enriched in MG63 (cluster 3), and 233 proteins were specific to HOS (cluster 4) (Figure S2B and Table S2A). The other clusters were enriched in proteins related to the experimental condition, with 194 proteins specific for NE condition (cluster 1), 197 proteins detected only for catTFREs C1 and C4 (cluster 5) and 288 proteins shared in both NE and catTFREs (cluster 6). While there were no TFs present in the NE cluster (cluster 1), clusters 2, 3, 4, 5 and 6 had many TFs. As expected, the top gene ontologies for biological processes of cluster 5, representing proteins pulled down by C1 and C4, were related to epigenetics, chromatin, and regulation of gene expression (Figure S2C). While for the other clusters, the top 5 biological processes were related to metabolic processes (cluster 1), intermediate filaments (cluster 2), extracellular structures (cluster 3), actin and RNA-related processes (cluster 4) and mitochondrial functions (cluster 6) (Figure S2C). Nuclear Factor I/B (NFIB) and RUNX2, two TFs known to promote tumour metastasis [41,42], were enriched by C1/C4 pulldown in MG63 (cluster 3) and HOS/U2OS (cluster 5), respectively (Figure 2B and Table S2A). TEA domain family member 1 (TEAD1), a transcription factor previously shown to promote proliferation in OS cells [43], was enriched in catTFREs C1 and C4 over NE in all three cell lines. Moreover, we found that JUNB, a TF and activator protein-1 (AP-1) complex component known to promote cell proliferation, was most highly enriched in C1 and C4 in MG63 (cluster 3). These four TFs were examples for enrichment by catTFREs C1 and C4.

**Figure 2.**
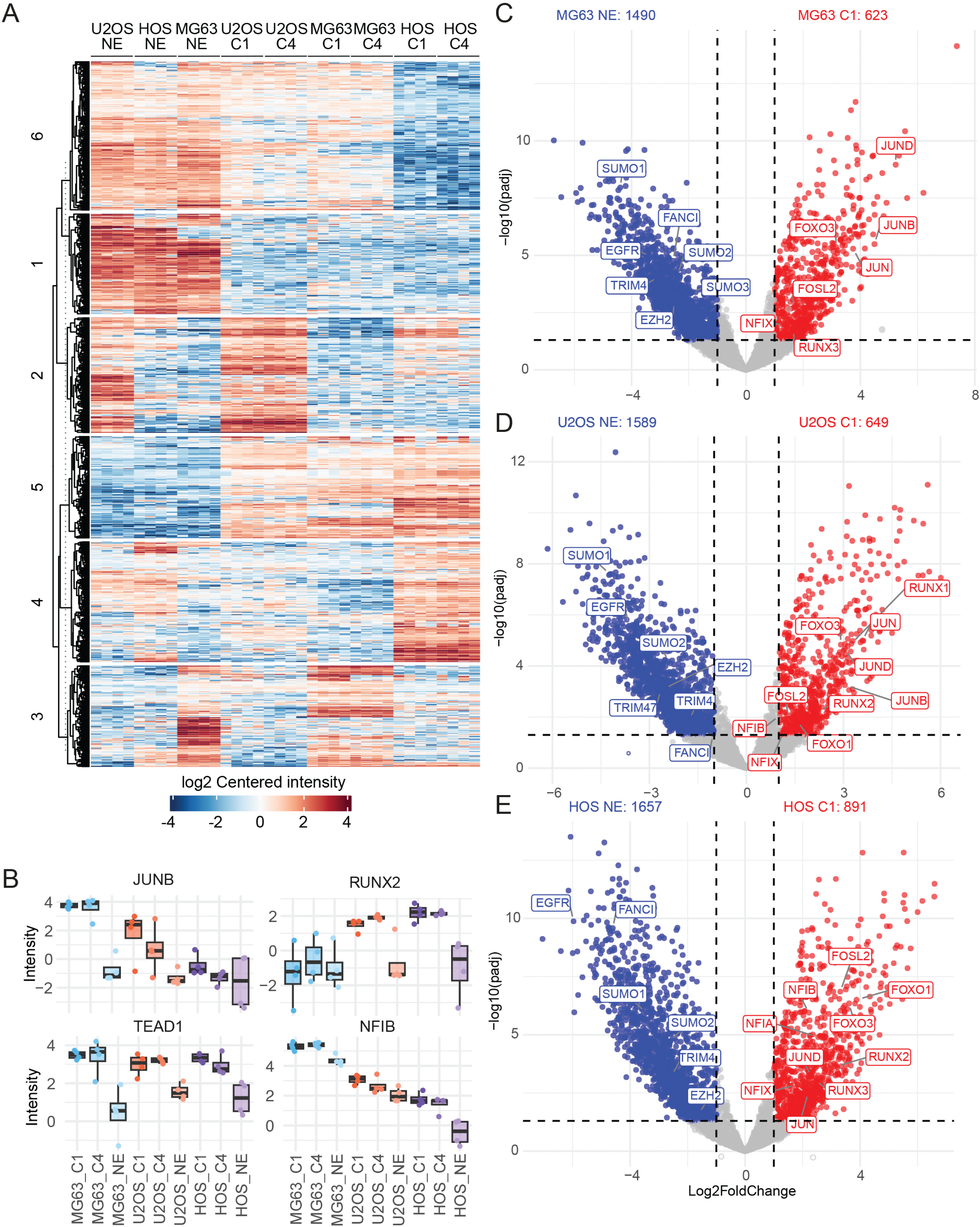
Differential protein analysis of the catTFRE approach versus nuclear extracts. A) Heat map of differential protein analysis between cell lines, C1, C4 and NE comparisons, defining clusters 1-6 (n = 4). B) Intensity levels of proteins JUNB and RUNX2 above; and TEAD1 and NFIB below, in NE and the catTFREs C1 and C4, for cell lines MG63, U2OS and HOS (n = 4). C, D, and E) Volcano plots of differential protein expression (DEP) analysis between C1 and NE in MG63 (in C), U2OS (in D), and HOS (in E), abs(log2FoldChange) > 1 and a p-value < 0.05. Significant proteins detected are coloured in blue (NE) and red (C1) with a selection of key proteins highlighted. All source data are provided in Table S2.

Proteins pulled down by C1 and C4 had relatively similar MS profiles and clustered together (Figure 1D). To further investigate the regulatory TF networks in different OS cell lines, we performed differential enrichment of protein (DEPs) analysis between the proteins pulled down by C1, C4 and proteins mapped in NEs. In all three cell lines, there were significant differences in protein enrichment between catTFREs C1, C4 and NE (Figures 2C-E, S2D-F and Table S2B-M). For MG63, a total of 2113 DEPs were detected in C1/NE (1490 up in NE and 623 up in C1), and 2269 DEPs were detected in C4/NE comparisons (1610 up in NE and 659 up in C4) (Figure 2C and S2D, Table S2B-C). Interestingly, only 32 and 40 TFs were enriched from MG63 NE, compared to 138 TFs pulled down with C1 and 92 TFs with C4. This included Fos related antigen 2 (FOSL2), JUN, JUNB, and JUND that are TFs that have been shown to play a role in osteosarcoma development and invasiveness [44–46] (Figures 2F and S2D). Similar results were observed for enrichment from HOS and U2OS cell lines when comparing DEP analysis between catTFREs and NE. A significant number of proteins were enriched in catTFREs of U2OS, with 2238 DEPs in C1/NE (1589 up in NE and 649 up in C1) and 2302 DEPs in C4/NE (1669 up in NE and 633 up in C4) (Figure 2D and S2E, Table S2F-I). This included a greater number of TFs (124 in C1 and 117 in C4 compared with TFs detected in U2OS NE (40 and 48). On the other hand, HOS catTFREs identified up to 2,548 DEPs in C1/NE (1,657 proteins up in NE and 891 in C1) and 2,556 DEPs in C4/NE (1,721 proteins up in NE and 835 in C4) (Figure 2E and S2F, Table S2J-K). We enriched 183 TFs C1 over 26 in NE, and 133 TFs in C4 over 35 in NE, respectively (Table S2L-M). Next, we compared how many of the TFs and coregulators specifically enriched by catTFRE C1 and C4 over NEs, that overlapped. We found that 93 TFs that overlapped between the three OS cell lines including all members of the TEAD family (TEAD1-4), Myocyte Enhancer Factor 2 (MEF2) family (MEF2A/C/D), ATF1-CREB1 complex, FOSL2, JUN and JUND (Figure S2G and Table S2N). Furthermore, we found 25, 16 and 46 TFs that were specific for each cell line. For example, PITX1 and Activating transcription factor 6 (ATF6) and Sine oculis homeobox 2 and 5 (SIX2/5) were enriched in MG63. When expressed PITX1 has a role in preventing OS metastasis and ATF6 act a marker of better prognosis [47,48], Amongst the TFs, we enriched RUNX1, TP53 and Signal transducers and activators of transcription 2 (STAT2) from U2OS and GATA4, SIX1 and STAT3 were enriched from HOS. SIX1 and STAT3 were linked to OS poor prognosis [49,50]. GATA4 is a regulator of RUNX2 in osteoblast bone mineralization [51]. There were 20 TFs enriched in both HOS and U2OS, including MYBL2, FOXO1, High mobility group AT-hook 2 (HMGA2), Special AT-rich sequence-binding protein 2) (SATB2) and RUNX2 were detected. All these TFs have previously been shown for promoting cell proliferation, and tumour immune pathways in osteosarcoma [52–54]. A total of 34 TFs were overlapping between HOS and MG63 with ALX3, RUNX3 and DNMT1 (Figure S2G and Table S2N).

The OS tumour epigenetic landscape states have recently been defined by TF footprinting analysis based on ATAC-seq DARs utilizing chromVAR [55] on a panel of OS PDX/PDX cell lines, osteoblast and OS cell lines [28]. MG63, HOS, and U2OS were found to be characterized by the late osteoblast-derived state (LOD) [28]. We therefore overlapped catTFRE enriched TFs from C1 and C4, with the TFs motif footprints for LOD and EOD state (Table S2O) [28]. Out of the 45 LOD TFs, we found six TFs (FOSL2, JUN, JUND, TEAD1,3 and 4) in all three cell lines and six in one or two cell lines: GATA4 (HOS), ZNF24 (U2OS), MAFG (MG63), JPD2 (HOS and MG63), JUNB and MAFF (U2OS and MG63). As these LOD and EOD transcriptional states were shown to coexist in OS tumours based on single cell analysis, we investigated whether any of the EOD TFs overlapped with our C1/C4 MS dataset. To our surprise we found that five out of 39 TFs common for all three cell lines (MEF2A, C and D, SP1 and SP3), and four TFs were enriched in one or two cell lines including RUNX1 (U2OS), RUNX2 (U2OS and HOS) and ALX and RUNX3 (MG63 and HOS). We have shown that by applying the catTFREs approach, we enrich more TFs than by direct MS analysis of polypeptides in NEs and that some of these TFs are specifically enriched in two or more of the OS cell lines.

### Osteosarcoma cell line specific transcription factor networks

We next assessed whether the TFs enriched by catTFREs from the OS cell lines can be used to identify the master regulatory TFs and networks involved in OS cancer progression and metastasis. U2OS and HOS cell lines exhibit higher metastatic abilities than MG63 [31] and we therefore sought to identify the differential protein enrichment of U2OS or HOS over MG63. To investigate the TF profiles associated with OS aggressiveness, we analysed the differential protein enrichment of C1 for each cell line. Principal component analysis and Pearson correlation showed that the MS data of the catTFREs C1 were separated between cell lines and correlated well between replicates (Figure S3A and B). There was a total of 868 DEPs between U2OS/MG63 (343 proteins up in MG63 catTFREs C1 and 525 proteins up in U2OS catTFREs C1), whereas up to 1165 DEPs were found between HOS/MG63 comparison (588 proteins up in MG63 catTFREs C1 and 577 proteins up in U2OS catTFREs C1) (Figure 3A and B and Table S3A and B). We found an overlap of 165 proteins enriched in MG63 in both conditions, including NFI and JUN protein families. In contrast, a total of 226 proteins were enriched in HOS and U2OS, highlighting several essential TF that might be responsible for regulating osteosarcoma metastasis, including HMGA2 [52], RUNX2 [56] and FOXO1 [39].

**Figure 3.**
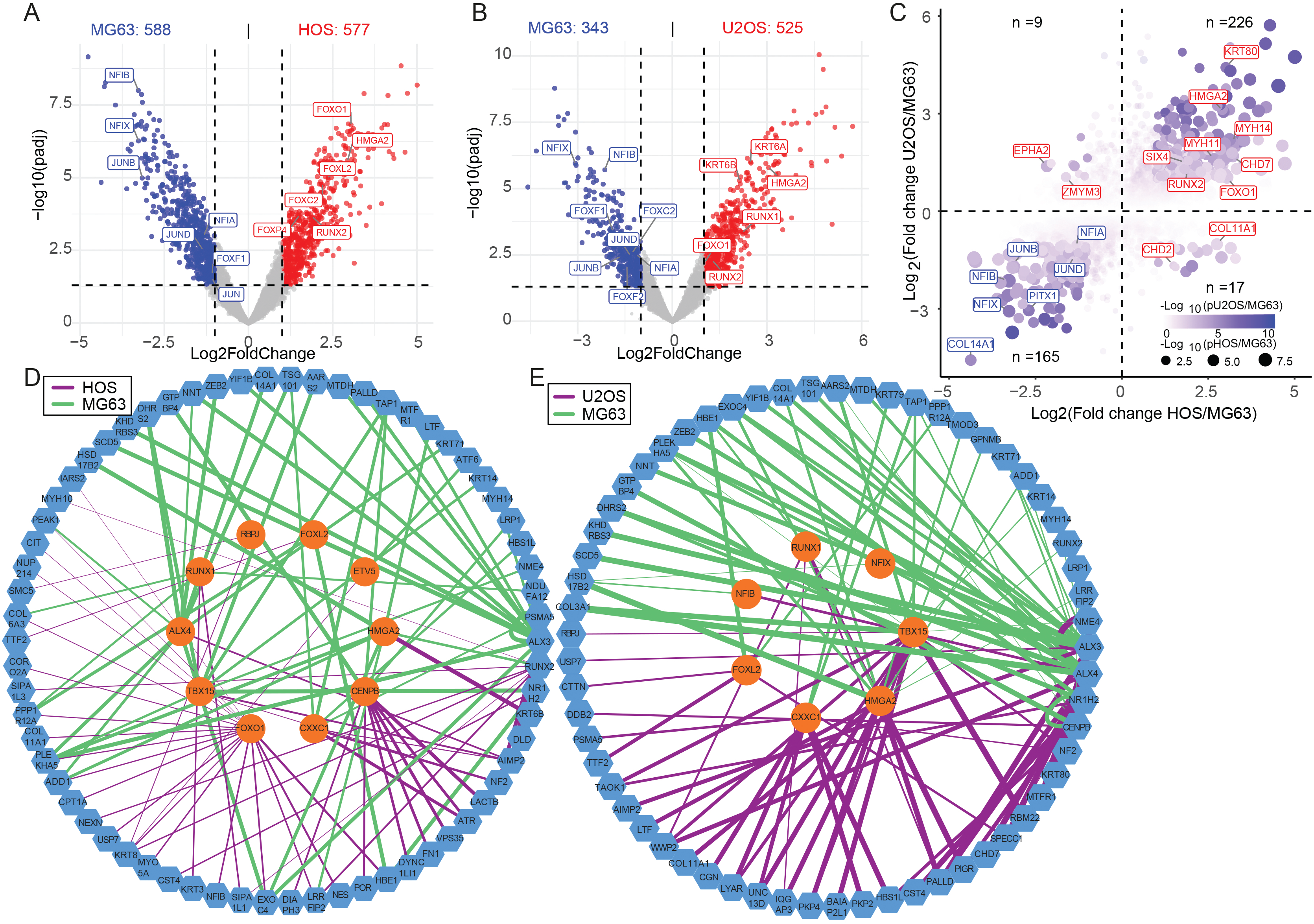
catTFREs C1 transcription factor networks. A) Volcano plots of the differential protein expression (DEPs) between catTFRE C1 for HOS (red) and MG63 (blue) cell lines. B) Volcano plots of the differential protein expression (DEPs) between catTFRE C1 for U2OS (red) and MG63 (blue) cell lines. C) Integration of the differential protein analysis for C1 from the HOS/MG63 and U2OS/MG63 comparison by log2Foldchanges and p-adjust values. Significant genes are clustered by log2FoldChange, highlighted by −log10(pValue HOS/MG63) and sized by −log10(pValue U2OS/MG63). D) Top 200 of the highest differential edge degrees between HOS/MG63, filtering based on the differential protein enrichment in A), plotted by Cytoscape. The edges in HOS and MG63 were highlighted in purple and green, respectively. E) Top 200 of the highest differential edge degrees between U2OS/MG63, filtering based on the differential protein enrichment in B). The edges in U2OS and MG63 were highlighted in purple and green, respectively. All source data are provided in Table S3.

We next utilized the Passing Attributes between Networks for Data Assimilation (PANDA) tool [57,58] to predict the specific protein-protein regulatory network between OS cell lines. We downloaded the prior motif-based TF-target gene (tool resources) and potential TF protein-protein interaction [59], and incorporated it with protein enrichment of the catTFREs C1 to predict the network models. The analysis weighted the edges between TFs and genes based on the protein enrichment C1 datasets and then modelled the aggregate regulatory network for each OS cell line. Compared to the differential networks analysis, we identified up to 54,877 specific edges between U2OS/MG63 (30,317 U2OS-specific edges and 24,560 MG63-specific edges) and 37,772 specific edges between HOS/MG63 (17124 HOS-specific edges and 20,648 MG63-specific edges) (Figure S3C and D). Subsequently, we filtered the TFs present in DEPs in U2OS/MG63 and HOS/MG63 (Figures 3A and 3B, Table S1E), and then ranked the nodes and edges based on their edge weights. The top 50 edges from each cell line from the comparison were selected to plot the PPI networks using Cytoscape [60]. The U2OS/MG63 and HOS/MG63 networks consisted of 72 nodes and 74 nodes, respectively, represented by different colours: MG63-specific edges were green, and HOS- or U2OS-specific edges were purple (Figures 3D and E). The scale of the edges illustrated the strong interaction between TFs and genes, and the shapes illustrated TFs (Ellipse, orange nodes) and genes (Hexagon, blue nodes). FOXL2, RUNX1, HMGA2, T-box transcription factor 15 (TBX15) and CXXC1, were the top orange node TFs in both networks. Patients with osteosarcoma have higher expression levels of HMGA2 and it has been linked to lower survival [52]. Nodes for HMGA2, CXXC1, TBX15 had edges with blue nodes in both U2OS and HOS cells whereas Centromere protein B (CENPB) and FOXO1 were specific to the HOS network. Moreover, we find that RUNX1 together with RUNX2 well known for having a role in osteogenesis [61] are represented by blue nodes interacting with HMGA2 in HOS and U2OS networks. These networks highlight the combinatorial action of TFs and chromatin regulators detected in MG63 and in both HOS and U2OS and in gene regulatory networks.

### Integration of catTFREs MS and RNA-seq reveals distinct commonalities between MG63 and U2OS versus HOS

To correlate gene expression with the catTFREs MS results, we performed RNA-seq from four biological replicates of MG63, HOS and U2OS. Principal component analysis and Pearson correlation profiles demonstrated that the RNA-seq data from each cell line were well-separated (Figure S4A and B). Next we compared the significant changes between U2OS/MG63 and HOS/MG63 to identify differentially expressed genes related to OS aggressiveness, (Figure 4A and B, Table S4A and B). The transcriptome comparison profiles were similar between U2OS/MG63 (3194 upregulated genes in MG63 and 4241 upregulated genes in U2OS) and HOS/MG63 (3490 upregulated genes in MG63 and 3588 upregulated genes in HOS). Common TFs up in HOS and U2OS over MG63 included *GATA4*, *RUNX2*, *SIX4*, *NFIA*, *HMGA2* and *MYB*. Amongst TFs up in U2OS were *GATA3* and *SRY-box transcription factor 2 (SOX2)* whereas *FOXO1, GATA2* and *Homeobox A1-3* (*HOXA1-3)* were up in HOS. We also compared gene expression changes between U2OS and HOS and identified 3910 upregulated genes in U2OS and 2584 upregulated genes in HOS (Figure S4C and Table S4C). To better characterize the differential gene expression across the three cell lines, we performed unsupervised clustering into five distinct subsets (R1-R5) (Figure S4D and Table S4D). Each cluster was represented by different TFs; for example, *RUNX2, TEAD4* and *FOXO1* were higher in U2OS and HOS (cluster R3) (Figure S4C, Table S4E and F). The expression levels of *VIM* and *Zinc finger E-box-binding homeobox 1 (ZEB1)* were up in MG63 (cluster R2), *JUN* was more prevalent in MG63 and HOS (cluster R1) and U2OS, and whereas *SOX4* and *SOX9,* a known osteoblast progenitor marker [62], were predominant in U2OS (cluster R4). *TWIST1,* a TF known to promote epithelial to mesenchymal transition, was expressed in both MG63 and HOS (cluster R5). When we conducted Gene Ontology (GO) analysis for significant gene expression changes between the OS cell lines, biological functions aligned closely between HOS and U2OS (Figure S4E). Unexpectedly for bone cancer development, common biological functions (HOS vs MG63 and U2OS vs MG63) relate to extracellular structures, renal development, neuronal development and axonogenesis. More expected was the common GOs linked to connective tissue development and skeletal system morphogenesis.

**Figure 4.**
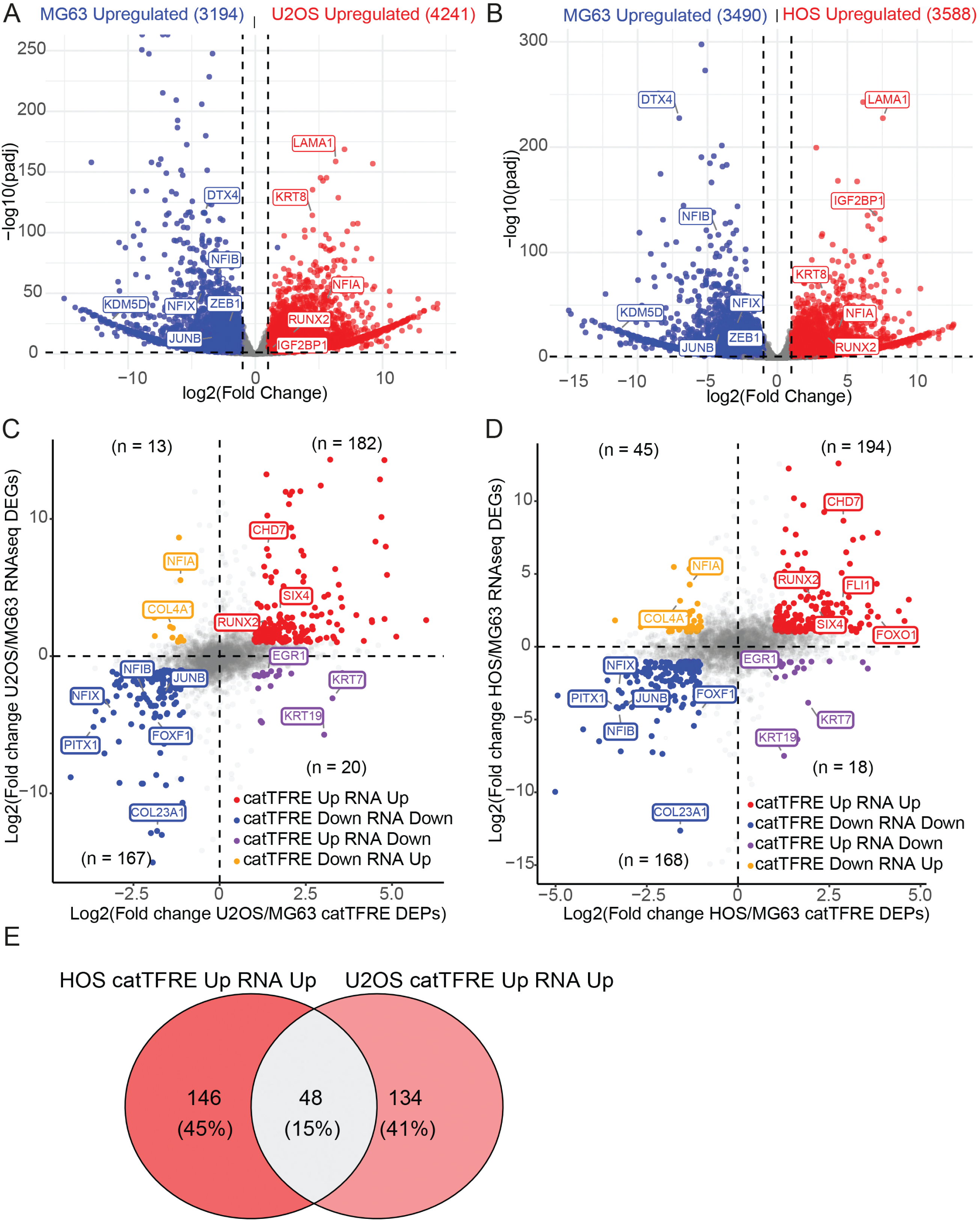
Integration of RNA-seq and catTFREs MS to identify transcription factor candidates that regulate OS malignancy. A) and B) Differential gene expression (DEGs) in the comparison of U2OS/MG63 and HOS/MG63. Statistically significant higher expression of HOS and U2OS genes are highlighted in red, while blue indicates higher expression of MG63. D) and E) Integration of the logFoldChange of DEPs and DEGs of U2OS/MG63 (in A) and HOS/MG63 (in B), with |log2FoldChange catTFREs C1| > 1 and |log2FoldChange RNA-seq| > 1. Four clusters are indicated by colours red, blue, purple, and yellow. Each cluster has a different number of proteins and key genes annotated. E) A Venn diagram of integrated datasets of U2OS and HOS for catTFRE up and RNA-seq up. All source data are provided in Table S4.

To further examine the correlation between catTFRE MS and RNA-seq, we integrated significant DEGs and DEPs (from the catTFRE construct C1) between HOS or U2OS versus MG63. We identified a subset of MG63 specific genes that were consistent in both the transcriptomic and protein levels (167 genes in HOS/MG63 and 168 genes in U2OS/MG63) (Figure 4C and D, Table S4E and F). TFs represented in the MG63 specific group included *NFIB, NFIX* and *JUNB*, which have been shown to suppress the metastasis process in cancer cells [42,63]. Furthermore, the integrated analysis revealed a correlation of upregulated genes for both DEGs with DEPs of U2OS/MG63 (182 U2OS-specific genes) and HOS/MG63 (194 HOS-specific genes). A total of 48 genes were common for catTFRE up/RNA-seq up in HOS and U2OS cell lines including *HMGA2, MYBL2, RUNX2*, *SIX4*, *CHD7*, and *FOXO1* (Figure 4E and Table S4G), which may be relevant in regulating the migratory ability of these two cell lines [39,64–66].

### Multi-omic analysis of Osteosarcoma epigenetic landscapes

We mapped the chromatin accessibility profiles by ATAC-seq [67] in MG63, U2OS and HOS for comparative global chromatin accessibility analysis, and integrated these with gene expression and C1 catTFRE proteomic data. The Pearson correlation and principal component analysis of global chromatin accessibility were shown to be distinct for the OS cell lines (Figure S5A and B). After identifying differential accessibility regions (DARs), we investigated the changes in opening regions between MG63 and U2OS or HOS. We identified 7,983 U2OS-specific peaks and 8,126 HOS-specific peaks of accessibility regions, followed by 7,816 and 8,159 MG63-specific peaks in the comparisons (Table S5A-C). Next, we sought to integrate ATAC-seq DARs at TSS spanning the TSS (+/− 500 bp) with RNA-seq and catTFRE C1 enriched proteins to cluster genes/proteins detected per cell line. When we compared HOS and U2OS with MG63, we identified 186 genes that showed significant changes in catTFREs C1 MS enrichment, RNA-seq, and ATAC-seq DARs at TSS regions, with a Pearson correlation coefficient greater than 0.5 in at least one pair of correlations. These were divided into four clusters where Cluster A (53 genes), Cluster D (44 genes), and Cluster C (36 genes) were cell line-specific for MG63, U2OS, and HOS, respectively (Figure 5A and Table S5E). Interestingly, cluster B (53 genes) presented a subset of proteins that are up in DEPs, DEGs, and DARs for both U2OS and HOS. We examined the correlation between proteins within each cluster by performing protein-protein interaction analysis using the StringDb database [59]. The protein-protein interaction networks that represented U2OS and HOS, cluster B, revealed a subset of TFs, including RUNX2, HMGA2, MYBL2 and FOXO1 (Figure 5B and Table S5E). When we performed GO enrichment analysis we found that cluster B was associated with terms including “chromatin”, “actin” and “cadherin” (Figure S5D and Table S5F). For cluster A, the protein network specific to MG63, many collagen family members (COL5A2, COL6A2, COL6A3, COL14A1 and COL23A1) were identified. Collagens are generally abundant proteins with different structural roles (Figure S5C, red colour) correlating with go terms associated with “extracellular matrix” for cluster A (Figure S5D). Extracellular matrix component remodelling is an intrinsic part of OS tumour progression [68]. We have demonstrated that the NFIB/X, PITX1, TBX15, JUNB/D TFs were pulled down from MG63 NEs (Figure 3 and Table S3), and this correlation was also supported by the integration of RNA-seq and DARs of the TSS region (Figures S5C, red colour). In the U2OS cluster C, we had enrichment of TFs PAX6 [69], and, HOXB4, ALX4 and Meis homeobox 2 (MEIS2), which correlate with go terms related to “core promoter sequence−specific DNA binding” and “minor groove DNA binding”. The U2OS-specific cluster also included nodes of KMT2A (MLL1), KDM2A (JHDM1A), which are epigenetic enzymes modulating histone 3 lysine 4 methylation, histone 3 lysine 36 demethylation (H3K36me2/1) respectively, whereas SENP3, is responsible for deSUMOylating of nuclear proteins such as TFs [70,71] (Figure S5C, turquoise colour). KDM2A binds unmethylated CpGs and contributes to the GO-term “unmethylated CpG binding” [72]. For the HOS cluster D (purple colour), the integrated analysis identified MYO1D, NES, NEFL and KRT family members which are associated with the GO-term “cell adhesion pathways” (Figure S5C). Taken together, integration of proteins enriched by C1 catTFREs, transcriptome and gene specific chromatin accessibility, gives us new knowledge on OS cell line-specific protein regulatory networks.

**Figure 5.**
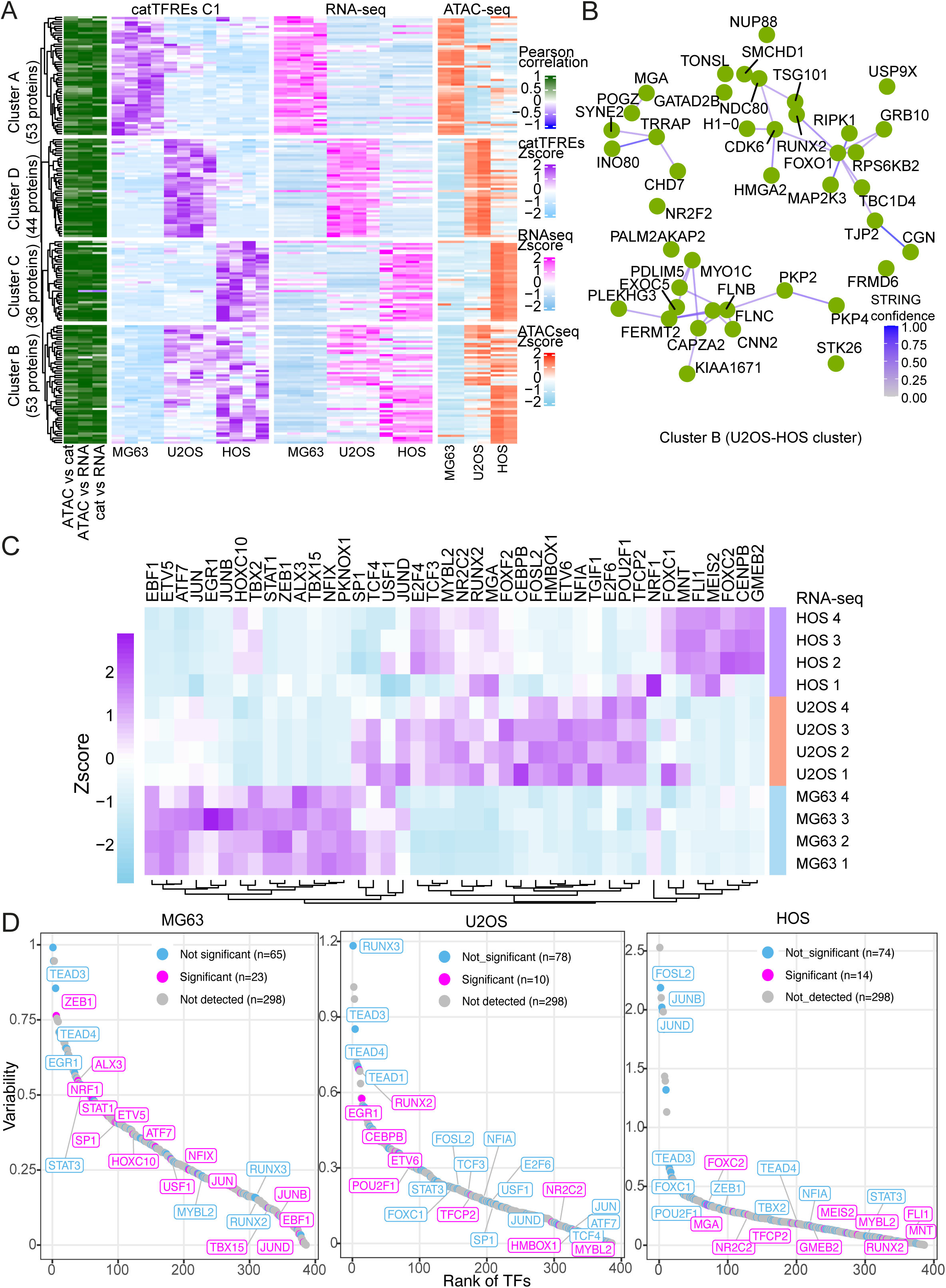
Overlapping catTFREs MS and transcription factor motif binding in chromatin accessibility. A) Heatmap visualizing the correlation of catTFREs C1 MS (Figure 3C), RNA-seq (Figure 4), and consensus peaks at TSS (ATAC-seq DARs) that are highly correlated. All genes/proteins were filtered based on the Pearson correlation between each pair of omics, where at least one pair had a correlation coefficient greater than 0.5. Four clusters are annotated where A is specific to MG63, B specific to HOS/U2OS, C specific to HOS and D specific to U2OS. B) Overview of high confidence of the predicted protein–protein interactions using the StringDb database. The cluster represents the protein and its interaction greater than 0.4 from the HOS/U2OS-shared cluster (Cluster B). C) Heatmap visualizing the normalized gene expression RPKM of all the TFs enriched in footprinting analysis in MG63, HOS and U2OS that were significant in catTFRE C1 comparison in Fig. 3C, 4C and D. Four replicates of RNA-seq per cell line. D) Ranking of TF footprinting from whole-genome consensus peaks in MG63 (left), HOS (middle) and U2OS (right) using chromVAR. TFs are coloured in blue (not significant), magenta (significant), and grey (not detected) for catTFREs C1 differential comparison from Fig. 3C, and Fig 4C and 4D integrated with RNA-seq, MG63/HOS or MG63/U2OS for each cell line. All source data are provided in Table S5.

We finally sought to identify the overlap between TF motifs from ATAC-seq footprinting analysis in all three OS cell lines, TFs enriched by catTFRE differential analysis with differential gene expression. The differential analysis allows us to identify TFs that are specific for each cell line but also common between HOS and U2OS as we are comparing C1 DEPs U2OS/MG63 and HOS/MG63 (Figure 3C) and C1 DEPs integrated with DEGs in RNA-seq (Figure 4C-D). Reads were systematically counted from the consensus peaks of the BAM files to ensure reliability and then computed across all OS cell lines to identify variability of TF motifs using chromVAR [55]. We detected footprints corresponding to 88 TFs, where 42 of these were significantly detected by the differential catTFRE C1 analysis (Table S5G). We plotted a heatmap to visualize gene expression levels of the 42 TFs and group these factors related to cell lines MG63, U2OS and HOS (Figure 5C). According to the heatmap all TFs were expressed and clustered. Of the 14 TFs more up in MG63, we find *ALX3, ETS variant transcription factor 5 (ETV5)*, *JUN, JUNB*, *NFIX* and *ZEB1*. The TFs *SP1*, and *TCF4* were more up in both MG63 and U2OS. Amongst the TFs more up in U2OS and HOS were *NR2C2, MYBL2* and *RUNX2*. Up in U2OS, were TFs such as *CEBPB, FOSL2* and *NFIA,* whereas HOS had upregulation of *FLI1, FOXC* and *MEIS2*. We repeated the analysis to distinguish the TF footprints that were significantly detected in C1 DEPs and DEGs for each individual cell line (Figure 5D and Table S5H-J). Highly ranked and C1 enriched TFs for MG63 were ETV5, JUNB, NFIX, and ZEB1. We enriched TFs, FLI1, FOXC2, MGA, and MEIS2 by C1 catTFRE in HOS and in U2OS we detected CEBPB, ETV6 and homeobox transcription factor 1 (HMBOX1). Common for both U2OS and HOS were significant detection of MYBL2 and RUNX2. We have identified subsets of TFs that are expressed, detected as proteins and represented as TF footprints in the chromatin landscapes that may reflect different invasion and migratory behaviours of the three different OS cell lines.

## DISCUSSION

TFs are considered relatively low-abundance proteins and are therefore more challenging to detect in proteomics. We have used the catTFREs approach [29] to enrich TFs and coregulators from three OS cell lines and captured a total of 7,594 proteins and 352 TFs across all conditions (Figures 1, S1 and Table S1). The MS analysis showed that we enriched more TFs in catTFREs C1 and C4 (310 and 269, respectively) compared to 244 TFs in NEs, across all cell lines (Figure 1B, Table S1F). Based on these results, we have shown that catTFREs approach can be used to enrich some TFs that may not be possible to map by MS directly from NE such as ALX3/4 and GATA4/6. However, when we compared TFs specifically enriched by catTFREs C1 and C4 over NEs, 38 % of overlapped in all three cell lines, and only 7%, 10% and 18% were specific to the individual cell lines (Figure S2G-H and Table S2N). Examples of cell line specific TFs enriched were ETV5 for MG63, GATA4 for HOS, NFKB1 for U2OS and FOXO1, HMGA2, MYBL2 and RUNX2 for HOS/U2OS. Based on this, we have shown that catTFRE can successfully enrich difficult to detect TFs and coregulators from 2D cell cultures. However, it may be of relevance to complement a catTFREs approach with MS analysis of total input proteins from NEs to identify the TFs that bind DNA and those with peptides detected in NE.

We have used cell lines as a model system to investigate whether OS aggressiveness is correlated with specific TFs and gene regulatory networks. MG63, HOS and U2OS used in this study represent different OS models (Table 1). For example, a deletion of *CDKN2A* has been associated with poor clinical outcome for OS, where MG63 and HOS carry homozygous deletion, and U2OS hemizygous deletion [73]. Only HOS carries a *TP53* missense mutation of arginine 156 to proline (R156P) that has been shown to promote migration and proliferation [73,74]. All three cell lines are highly proliferative, however we have focused on identifying TFs that may promote HOS and U2OS characteristics of higher invasive and migration ability [31].

In the least aggressive OS cell line, MG63, the proteomics reported specific proteins, including PITX, and members of the NFI and JUN families (Figure 2C, S2D, 3, S3E, Table S2B-E and S3). Interestingly, PITX1 is highly expressed in MG63 by the human cell atlas [75] and when overexpressed has been reported to prevent OS cell proliferation and migration [48]. TFs JUNB and JUND can both prevent metastasis while also being oncogenic [63,76,77]. To identify potential TFs related to the higher migratory and invasive potential of U2OS and HOS, we performed differential analysis with MG63 C1 catTFRE. Although carrying different mutations with and more than 500 proteins showing cell line-specific expression, HOS and U2OS cells exhibited enrichment of many similar TFs, including HMGA2, RUNX2, SIX4 and FOXO1 (Figure 3 and Tables S2F-M and 3). The upregulation of key regulators of osteoblasts, including RUNX2, HMGA2, CDK6, and FOXO1 [37,78–80], have been shown to promote OS progression and metastasis [38,39,81]. Our PANDA prediction of protein-protein interaction network illustrated the top candidates of transcriptional regulators in the OS cell lines. Amongst most interacting candidates that regulate transcriptional expression in HOS and U2OS were FOXO1 with 20 and 8 edges of connections, while RUNX2 with 15 and 5 edges of connections (Figures 3D and E). FOXO1 is a master TF [82] and RUNX2 regulate chromatin in the osteoblast lineage [37] suggesting these are transcriptional drivers of OS.

Despite well-developed analyses of MS, an omics integration was necessary to better characterize the OS transcriptional regulatory networks. Our multi-omics analysis revealed a high correlation between proteomics, transcriptomics, and chromatin accessibility. Approximately one-third of catTFREs C1 DEPs overlapped with RNA-seq DEGs (182/525 U2OS DEPs, 194/577 HOS DEPs, and 168/588 MG63 DEPs) (Figure 4). We then filtered the overlaps of catTFREs MS, RNA-seq, and ATAC-seq based on each pair of correlation analyses, which helped us identify a specific set of genes where changes in MS data also correspond to changes in gene expression and chromatin accessibility. The four sub-clusters of genes showed differences and similarities between the OS cell lines, with three OS cell line-specific genes (cluster A, C, and D) and one cluster that was shared by U2OS and HOS cell lines (cluster B) (Figure 5A-B and S5C). In the STRING networks, we observed a diversity of connections in the four clusters. For example, RUNX2 and HMGA2 were linked with CDK6 in the HOS/U2OS cluster. A clinical trial for CDK4/CDK6 inhibitor palbociclib in advanced sarcoma has shown promise with a close to 8 months progression-free survival in Osteosarcoma patients [83]. Furthermore, two chromatin remodelling factors CHD7 and INO80, known to modulate chromatin and promote transcriptional expression of oncogenes [65,84,85], clustered with Transformation/transcription domain-associated protein TRRAP which when overexpressed correlate with poor OS prognosis [86]. In the MG63 cluster A, TFs NFIB and NFIX were connected with PSIP1. PSIP1 has been shown to promote genome integrity by reduction of R-loops and promote cancer proliferation [87]. There were also many collagen proteins enriched in the MG63 specific cluster. We detected SP1 by the catTFRE approach in MG63 (Table S3B) and SP1 has also been shown to promote cell proliferation and regulate expression of different collagen types in OS [25,88].

When we performed a multi-omics integration of catTFREs C1 TFs and coregulators with RNA-seq and ATAC-seq footprints, we detected 88 TF motifs overlapping catTFREs data, where 42 of these TFs were significant based on differential comparisons (DEP/DEGs) of HOS/U2OS over MG63 (Figures 3C, 4C-D). We represented the 42 TFs in a gene expression heatmap and a footprint ranking per cell line (Figures 5C and D and Tables S5G-J). The transcriptional regulation process in lineage commitment relies on the pioneering TFs that bind DNA on nucleosomes and open chromatin for other TFs [89–92]. Common factors differentially enriched in HOS and U2OS (catTFRE Up RNA Up Figure 4D) were RUNX2 and MYBL2. RUNX and MYB family members have been shown to have pioneering abilities [37,93–95] and MYBL2 and RUNX2 are implicated in OS [9,11,21,53,56]. These TFs were enriched by catTFRE in U2OS and HOS and may likely be involved in promoting the cell lines’ aggressiveness phenotype. In U2OS, TFs such as CEBPB, ETV6 and HMBOX1 were highly ranked, all up in RNA and catTFRE (catTFRE Up RNA Up Figure 4D). Both CEBPB and HMBOX1 have been shown to have inhibitory effect on OS cell proliferation and metastasis [96,97] and may therefore not be the major players in U2OS aggressiveness phenotype. On the other hand, ETV6-NTRK3 fusion protein promotes proliferation and counteracts ETW-FLI1 fusion protein in Ewing sarcoma and ETV6-NTRK3 was also a driver in a rare type of extraskeletal OS [98,99] suggesting that ETV6 may be a driver in U2OS. Among the significant footprints in HOS, we detected FLI1, FOXC2, and MEIS2 (catTFRE Up RNA Up, Figure 4D). The FLI1-EWS fusion has the FLI1 C-terminal ETS DNA binding domain and is an oncogene that remodels the chromatin landscape in Ewing sarcoma [100]. Interestingly, an OS model with EWS-FLI1 prevented osteogenesis and promoted OS development [101]. Furthermore, FOXC2 (catTFRE Up RNA Up, Figure 4D) is a driver of osteoblast development and OS invasiveness and migration abilities [102,103]. Similarly, MEIS2 was upregulated in metastatic OS [104]. Early growth response 1 (EGR1) footprints were ranked as number 14 in U2OS, 108 in HOS and 311 in MG63, however the gene expression was lower in U2OS/HOS than MG63 (catTFRE Up RNA Down, Figure 4D). Interestingly, ectopic overexpression of EGR1 in HOS has previously been shown to prevent migration but had no effect on proliferation [105]. ZEB1 was significant and the sixth highest ranked footprints in MG63. ZEB1 promotes epithelial to mesenchymal transition and upregulation in OS cell lines showed dual roles increasing colony formation in MG63 and promoting metastasis of cell line 143B [106]. MG63 also had significant enrichment of ALX3, Activating Transcription Factor 7 (ATF7), ETV5, JUNB, NFIX, and STAT1. ETV5 is upregulated in many cancer subtypes and a study showed that high expression in SaOS-2 cells increased proliferation and invasiveness [107,108]. ATF7 and JUNB are part of the activator protein-1 (AP-1) complex known to promote cell proliferation and OS tumour invasiveness [45,76]. However, increased levels of NFIX and JUNB/D have negative effects on cancer metastasis [42,63]. The less aggressive MG63 phenotype may also be promoted by STAT1 as it highly upregulated (Figure 5C and Table S4A). Moreover, SP1 was as a suppressor of metastasis when upregulated in OS cell lines [109,110].

Heterogenous OS tumours were recently suggested to be divided into two epigenetic early and late osteoblast states (EOD and LOD) defined by chromatin accessibility analysis [28]. Based on this study [28], MG63, HOS and U2OS cell lines were all grouped within the LOD state having chromatin accessibility regions enriched in TF motifs including FOSL and JUN TF families. Sweet-Cordero and colleagues integrated ATAC-seq with RNA-seq, however they did not map TFs by proteomics. Among the 244 TFs and co-factors enriched by catTFRE C1 and C4 over NE, we identified 12 of the LOD factors by catTFRE proteomics (27% of 45 LOD TFs). Moreover, we also detected nine EOD factors (23% of 39 EOD TFs) in the three cell lines (Figures S2G-H). Based on our catTFRE proteomics, we were not able to show that MG63, U2OS and HOS belonged specifically to LOD nor EOD, rather we enriched TFs of both states. In our differential proteomic analysis, the presence of the FOS/JUN family was more present in MG63 suggesting a LOD state, however the presence of MEF2AB and C in the same cell line also show that it could be in an EOD state (Figures 2C and 3C). Our study relies on a proteomics integrational analysis with gene expression and chromatin accessibility based on a very high sequencing dept, where catTFRE has the advantage of sensitively detecting TFs present in the OS cell lines. An ongoing question is what TF protein expression levels are necessary to stimulate gene expression effects and even cellular reprogramming, as well as whether some TFs act alone or in concert with other TFs. TFs FOXA1 and HNF4A that can act together or alone in activating liver and intestine genes [111]. A recent study has shown that the expression of TFs including TWIST1, JUN and STAT3 changes in OS cell lines upon metastatic progression in mice and that this is correlated with dynamic change in the epigenetic landscape [112]. Moreover, evolutionary OS intratumour studies have shown that although these heterogenous mesenchymal malignancies are genomically homogenous, the genomic structural rearrangements take place relatively early in tumorigenesis [113]. Biological characteristics of heterogeneous OS tumour development, invasion, metastasis and drug resistance are therefore more likely to be regulated by TFs and dynamic changes in the epigenetic landscapes. We suggest that more comprehensive analysis is needed to define the TFs that are actively taking part in LOD and EOD states, and that there may be more dynamic epigenetic substates in OS.

Here, we have shown that our catTFRE approach has enriched TFs and an interacting sub-proteome from MG63, HOS and U2OS OS cancer models. We have identified TFs and coregulators specific for each cell line and common for HOS and U2OS. Combining catTFRE TF enrichment strengthens chromatin accessibility analyses. Taken together, our integrative multi-omics analysis of the transcription regulatory network provides new understanding of the complex regulation of highly malignant bone tumours and offers rationales for refinement of epigenetic subtypes and therapeutic targets.

### Limitations of the study

In this study, we have used a catTFREs pull-down approach with two different biotinylated 3 kilobase pair long DNA sequences, C1 and C4, and compared the protein enrichment with MS detection in NEs from three different OS cell lines. Our focus was on three well characterized 2D cultured cell lines representing different genetics, and aggressiveness however we find commonalities and differences regarding epigenetic regulation. OS is heterogenous tumours and expanding the datasets with more models would strengthen the finding of this study. Although C1 was designed to have duplicates of 110 TFBS, the CiiiDER analysis on C1 detected a total of 632 TFBS. We also applied a control DNA, C4, digested from a reporter vector that contains fewer TFBS, and the CiiiDER analysis confirmed this by detecting 396 TFBS (Figure S1B and Table S2). There was an overlap of 351 TFBS between C1 and C4, however a total of 287 of the TFBS were present 10 or more times in C1 compared to only 62 of the TFBS in C4 being present 10 or more times. This was also reflected in that the number of TFs enriched in the pulldown was higher for C1 than C4 (Figure 1B). However, the enrichment of TFs on the DNA did not correlate well with specific TFBS design nor CiiiDER prediction (Figures S1A-D). This may be explained by the fact that DNA binding affinity varies between different eukaryotic transcription factor families, as well as a TFs ability to bind sequence motifs that are not identical with different affinity [114]. Different models on mechanisms of TFs DNA binding have been proposed, and approximately 90% TFs bind to DNA at any time in the nucleus with a high on off rate, suggesting that TFs bind both non-specifically and specifically to DNA [115]. Moreover, the residency time of a specific TF on DNA may vary. It is likely that the TFs move by sliding or hopping on the DNA until the TF identifies its respective TFBS [116]. As catTFRE pulldown is an in vitro assay, TFs residency time is beyond the scope of this study.

When performing StringDB analysis of confidence predicted protein - protein interactions, keratin family members were prominent in the HOS specific cluster (Figure S5C, purple). From a MS perspective, these proteins may be perceived as contaminants of the preparation. However, several keratin genes were significantly upregulated in OS cells in RNA-seq (Figure 3, 4 and Table 3) and detected as DARs (Table S5) and were therefore considered highly relevant for OS regulation in our multi-omics analysis. Different keratins have in recent years been found to be both prognostic markers and drivers of cancer progression in different types of cancer [117]. In OS, KRT17 is highly expressed and has been shown to promote cell proliferation [118]. Also, KRT19 has been suggested to promote OS tumour invasion via deregulation of lysosomal-associated membrane protein 3 [119]. Moreover, KRT80 was identified as upregulated in OS cell lines and deregulated by a BRD4 inhibitor (GNE-987) through deregulation of histone H3K27 acetylation at the gene locus [120]. In cutaneous squamous cell carcinoma, co-expression of KRT8 and KRT18 has been shown to increase metastasis and invasiveness [121]. Taken together, aberrant transcriptional regulation of different keratins may contribute to the aggressiveness of HOS and U2OS.

A significant proportion of proteins pulled down by the catTFREs C1 and C4 were not DNA-binding proteins (Figure 1B). Among these proteins we identify TF co-factors, co-regulators, DNA repair factors, other chromatin associated proteins but also extracellular matrix factors and components of the cytoskeleton that may be non-specific protein interactions. Further studies are required to optimize the conditions to reduce non-specific interacting proteins. Increasing the number of TFBS per TF or reducing the DNA length may improve the specific enrichment. Further optimizations will enable us to upgrade the catTFRE approach to enrich more TFs and their coregulators.

## METHODS

### Cell culture

Human osteosarcoma HOS, U2OS, and MG63 cell lines (CRL-1543, HTB-96 and CRL-1427, Table 1.) were purchased from the American Type Culture Collection (ATCC). U2OS cells were cultured in DMEM media (ThermoFisher #41965039) supplemented with 10% foetal bovine serum(FBS) (Capricon Scientific, #FBS-11A) and 1% penicillin and streptavidin (P/S) (GibcoTM #15140122). HOS and MG63 cells required the addition of 1 mM sodium pyruvate (Gibco, #11360070) to the DMEM media containing 10% FBS and 1% P/S. Cells were grown at 37 °C and 5% CO2, passaged every 72 hours with fresh medium, and regularly checked for mycoplasma.

### Nuclear protein extraction

The cells were harvested by trypsinization, washed twice in 1x phosphate buffered saline (PBS) and pelleted at 182 rcf for 10 minutes. The cells were resuspended in hypotonic buffer (10 mM 4-(2-hydroxyethyl)-1-piperazineethanesulfonic acid (HEPES) pH 7.5, 2 mM MgCl2, 25 mM KCl, 1X cOmplete Protease inhibitor cocktail (Roche), 0.2 μM Phenylmethylsulfonyl fluoride (PMSF) and 1 mM dithiothreitol (DTT)) on ice for 20-40 minutes depending on cell line. To release the nuclei, douncing was performed with a 7 ml glass douncer with B (tight fit) pestle on ice and a small sample was monitored with a phase contrast microscope until most of the nuclei were released. Nuclei were counted and sucrose was added to a final concentration of 0.25 M and the sample mixed by inversion. The supernatant was removed and the nuclei were washed once in buffer N (10 mM HEPES pH 7.5, 2 mM MgCl2, 25 mM KCl, add fresh: 250 mM sucrose, 1mM DTT, 1mM PMSF) before being resuspended in buffer BC100 (25 mM HEPES pH 7.5, 1 mM, MgCl2, 0.5 mM EGTA, 0.5 mM EDTA, 100 mM NaCl, 10% Glycerol, add fresh: 1mM DTT, 0.2 mM PMSF, 1x Protein inhibitors (PI)), to a final concentration of 50 000-150 000 nuclei/μl. Nuclei were incubated on ice for 30 minutes and then sonicated with a Biorupter Pico (Diagenode) for 5 cycles (30 sec on/off, high setting) at 4 °C. Debris was removed by centrifugation at max speed for 10 minutes at 4 °C. The nuclear extract (NE, supernatant) was flash-frozen on dry ice and transferred to a −80 °C freezer for downstream experiments. If not used immediately, the NEs were stored at −80 °C and gently thawed and centrifuged at max speed before caTFRE pulldown.

### catTFREs protein pulldown and MS protein analysis

The catTFREs C1 was designed with a 3274 base pair insert of a total of 110 transcription factor binding sites (TFBS) in duplicates separated by three nucleotide linkers and synthesized and cloned into pUC57 using SalI-PstI by Genscript. As control, a sequence from pGL4.26[luc2/minP/Hygro] plasmid (Promega) that has been designed to have a reduced number of TFBS, was used. A site for PvuII was removed in the luc2 gene in pGL4.26 by the QuikChange® Site-Directed Mutagenesis Kit (Stratagene) resulting in Q42H mutation (pGL4.26PI-). The 3 kilobase inserts were linearized. The vector containing C1 was digested with restriction enzymes Eco53KI and BamHI (New England Biolabs). Vector pGL4.26PI- was digested with restriction enzymes PvuII and BamHI (New England Biolabs). The restriction digestion reactions were precipitated and biotinylated by biotinylated dUTP and dATP (Roche) using a Klenow Fragment (New England Biolabs). The Klenow enzyme was inactivated for 20 minutes at 70 °C and removed together with nucleotides by centrifugation over Sepharose G50 columns (Roche). The vector backbones were separated from biotinylated C1 and C4 with a restriction digest with PstI and confirmed by agarose gel electrophoresis before downstream applications. The biotinylated TFBS fragments were immobilized on DynabeadsTM M-280 Streptavidin (Invitrogen, 11206D) by incubation for 3 hours in KilobaseBINDER (Invitrogen, 60101) on a rotator at 4 °C following the manufacturer’s instructions. Dynabeads with C1 or C4 were washed three times with Wash Solution (10 mM TrisHCl pH 7.5, 1 mM EDTA, 2 M NaCl) to remove nonspecific and unbound DNA and once with TE buffer (10mM TrisHCl pH 7.5, 1mM EDTA). The C1 and C4 Dynabeads were blocked with two washes of 1x PBS containing 0.5% Bovine Serum Albumin (BSA). The beads were resuspended in BC100 buffer (25 mM HEPES pH 7.5, 1 mM MgCl2, 0.5 mM EGTA, 0.5 mM EDTA, 100 mM NaCl, 10% Glycerol,1X cOmplete Protease inhibitor cocktail (Roche), 0.2 μM PMSF and 1mM DTT).

The nuclear protein extracts from either MG63, U2OS or HOS were added at a ratio of 100 μg protein to 5 μg linearized DNA on Dynabeads in the BC100 buffer. The pull-down reactions were incubated by rotation for 2 hours at 4 °C, then washed three times with BC100 buffer, two times in 1xPBS + 0.01%NP40 and eluted in 20 mM ammonium bicarbonate for MS analysis.

Samples were analysed by LS-MC timsTOF Pro (Bruker Daltonik, Germany) and searched against the human Uniprot database using PEAKS X+ version 10 (Bioinformatics Solutions, Canada). The following parameters were applied: digestion enzyme: trypsin; maximum missed cleavage: 1; fragment ion mass error tolerance: 0.03 Da; parent ion error tolerance: 15.0 ppm; fixed modifications: carbamidomethylation, maximum number of PTMs: 2, and false-discovery rate: 1%.

All protein and peptides matching with databases were counted and analysed by R (v4.5)/Bioconductor (v3.20) using the package DEP (v1.31.0). Samples identified in all replicates of at least one condition were filtered (thread = 0) and then processed for normalization and imputation of missing data using random draws following a Gaussian distribution (fun=“MinProb”, q=0.01). The data were then analysed for the Differentially Expressed Proteins (DEPs) among the different tested conditions, which were manually identified based on the aggressiveness of the osteosarcoma cell lines (HOS/MG63 and U2OS/MG63). Volcano and heatmap plots were visualized by the R/Bioconductor package ggplot2 [122].

### catTFREs WGCNA and PANDA regulatory networks analysis

All proteins were identified by MS from 36 samples (3 cell lines, 4 replicates, 3 experimental catTFREs C1, C4, and NE) using as input for WGCNA [123]. We mapped the MS data against the UniProt protein database, and the raw read counts were then analysed using the DEP R package [124]. Data were calculated for the scale of the topology model fit at 90% (h=0.9) and chose the power nearest the threshold (picked_power=7). Traits were assigned to experimental conditions (cell lines and catTFREs C1, C4, and NE) to classify subsets of proteins that were expressed similarly. The analysis was conducted using the unsigned consensus network with the following parameters: TOMType=“unsigned”, networkType=“signed”, mergeCutHeight = 0.15, and minModuleSize=100; the rest of the steps were set to default. Modules were aligned to colours, and then a heatmap was plotted with the correlation values for the traits.

MS spectral counts from C1 were used to build the gene regulatory networks. TF human motifs (Homo sapiens, hg38) were downloaded from PANDA resources (https://sites.google.com/a/channing.harvard.edu/kimberlyglass/tools/resources, accessed July 10, 2023), and the protein-protein interaction network was retrieved from the StringDb database [125] (https://string-db.org, accessed July 10, 2023). We utilized PANDA from *netZooR* [126] to establish the gene regulatory network, based on protein spectral counts, protein-protein interactions, and TF motifs with standard parameters. PANDA can predict and compare the weights of each protein-protein interaction between cell lines, identifying cell line-specific network edges. Results can be used for Gene Set Enrichment Analysis to plot the top 5 and the Enrichment plot of the GO:BP pathway. The transcription regulatory network was then filtered by a subset of the significant DEPs in the catTFRE C1 of HOS/MG63 and U2OS/MG63, to visualize the specific network between the comparisons using Cytoscape v3.10.2 [60].

### RNA-seq preparation and bioinformatic analysis

RNA was extracted from the cells using TRIzol (ThermoFisher, 15596018) and following the RNeasy Mini Kit (QIAGEN, 74106). In brief, cells were cultured in a 6-well plate, quickly washed with cold PBS on ice, and then lysed with 700 μL of Trizol. The homogenized lysate of the cell was incubated at RT for 5 min before being vigorously mixed with 140 μL of chloroform. After a short incubation at RT, samples were then centrifuged at high speed for 15 minutes, and the upper aqueous phase was transferred to new tubes for RNA isolation. A 1.5x volume of 80 % Ethanol was added, and RNA was extracted using the RNeasy Mini Kit according to the manufacturer’s instructions. Samples were measured for concentration and quality before being sent to the Novogene Europe, Cambrige, UK for library prep with polyA enrichment and sequencing. Transcriptomic data was acquired by Illumina NovaSeq 6000 sequencer using 150 bp paired-end settings.

For the bioinformatic analysis, fastq files were trimmed off adapters by the fastp (v0.23.4) tool [127] using paired-end input. The trimmed fastq files were then aligned with hg38/gencode v47 indexing using STAR alignment (v2.9.2) tool [128] with standard parameters and excluding the multimap reads. Raw reads were counted using featureCounts (Subread v2.0.4) tool [129], and differential gene expression was analysed using DESeq2 [130], following the guidelines. Gene Ontology analysis for each DEGs was performed using the *clusterProfile R* package [131] with standard parameters.

### ATAC-seq preparation, bioinformatic analysis and integration with omics

ATAC-seq samples were processed using the ATAC-Kit (Active Motif, #53150), which has been well described previously (Corces et al., 2017; Grandi et al., 2022). In brief, OS cells were prepared from 2 biological replicates then harvested by trypsinization, washed with cold PBS, and counted before proceeding with the ATAC library preparation. Approximately 100,000 cells were lysed in 100 μL of lysis buffer, followed by the pelleting of nuclei. The samples were then subjected to tagmentation using mutant transposase Tn5 for 30 minutes, and then DNA purification was performed. Amplification of the samples was carried out via a low-cycle PCR using an indexed primer set to prepare the library. For the final step, ATAC-seq samples were purified using SPRI beads in preparation for sequencing. Samples were sequenced with a sequencing depth of at least 200 million reads using Illumina HiSeq X.

Fastq files were processed to remove adapters and low-quality reads using *fastp* (v0.23.4) tool [127] with the default paired-end option. After trimming, reads were aligned with the human hg38 reference by *bowtie2* (v2.5.1) tool [132] with parameters: “*-X 2000 --threads 30 --end-to-end --very-sensitive*”. BAM files were maintained only for correctly mapped pairs (*-f 2*), and the unmapped or duplicated reads (*-F 1548*), and reads from mitochondrial or nonprimary chromosomes were excluded using *SAMtools* (v.1.18) [133] and MarkDuplicates (picard v3.0.0, https://github.com/broadinstitute/picard). *MACS2* [134] then called peaks using the following parameters: “*-nomodel -shift 100 -extsize 200*”. To define reproducible peaks between replicates, *BEDtools* (v2.28.0) intersect was used to identify a reciprocal overlap threshold of 50% of peaks with options: “-f 0.50 -r -u”. Peaks that overlapped with the blacklist (Amemiya et al., 2019) were excluded from downstream analysis. Subsequently, ATAC peaks were merged across all cell lines and replicates then *featureCounts* (*subread* v2.0.2) was quantified consensus peaks, and *chromVAR* (v1.30.1) tool [55] was used to identify the TF motif activities, which were then integrated with DEPs. The normalized peaks of the transcription start site (+/− 500 bp from TSS) consensus peaks were counted using the *bigWigAverageOverBed* tool [135], filtered to include peaks with a mean0 value of at least 20 over the background level. Differential accessibility regions (DARs) were then analyzed using *DESeq2* and the overlap of DEPs, DARs, and TF motifs were plotted with *ggplot2* [122]. *clusterProfile R* [131] and *STRINGdb* [125] packages facilitated the GO term analysis and enhanced visualization of interactions within each cluster derived from the integration results.

## CREDIT AUTHORSHIP STATEMENT

Conceptualization, R.E. and B.T.; Methodology, E.L.B.M., X.T.N., M.A., M.R., T.T.T, R.E. and B.T; Investigation, E.L.B.M., X.T.N., M.A., M.L., T.T.T.; Formal analysis, XTN; Visualization, X.T.N. and R.E.; Writing-original draft, X.T.N. and R.E.; Writing-review & editing, E.L.B.M., X.T.N., M.A., M.L., T.T.T., M.R., B.T. and R.E; Funding acquisition, R.E.; Resources, R.E. and B.T.; Supervision, M.R., B.T. and R.E.

## DATA AND CODE AVAILABILITY

RNA-seq data have been deposited at GEO with the accession number GSE308074 and ATAC-seq data has the accession number GSE309682. The MS proteomics data have been deposited to the ProteomeXchange Consortium via the PRIDE [136] partner repository with the dataset identifier PXD070562, and will be publicly available from the date of publication. The code has been deposited at EskelandLab Github/OScatTFRE. Any additional information required to reanalyse the data reported in this paper is available from the lead contact upon request.

## ACKNOWLEDGMENTS

This work was supported by the Norwegian Centres of Excellence scheme grant 262652 - CanCell (R.E.). The sequencing service was provided by the Norwegian Sequencing Centre (www.sequencing.uio.no), a national technology platform hosted by the University of Oslo and Oslo University Hospital, and supported by the Research Council of Norway and the Southeastern Regional Health Authorities. The informatic analysis was performed at Saga super computing resources (Project NN9632K) provided by UNINETT Sigma2 - the National Infrastructure for High Performance Computing and Data Storage in Norway. We thank Christian Koehler (Department of Biosciences, University of Oslo) for technical support, Anthony Mathelier (Norwegian Centre for Molecular Biosciences and Medicine, University of Oslo) for providing extracted TF binding motifs from JASPAR database and Ankush Sharma (Institute of Clinical Medicine, University of Oslo and Oslo University Hospital) for advice on bioinformatic analysis.

## DECLARATION OF GENERATIVE AI AND AI-ASSISTED TECHNOLOGIES

During the preparation of this work, the author(s) used Grammarly and Wordtune to correct the English grammar. After using these tools, all author(s) reviewed and edited the content as needed and took full responsibility for the content of the publication.

