## Supplemental figures for "Proteome-wide multi-omics profiling of osteosarcoma transcription factor networks"

### SUPPLEMENTAL INFORMATION

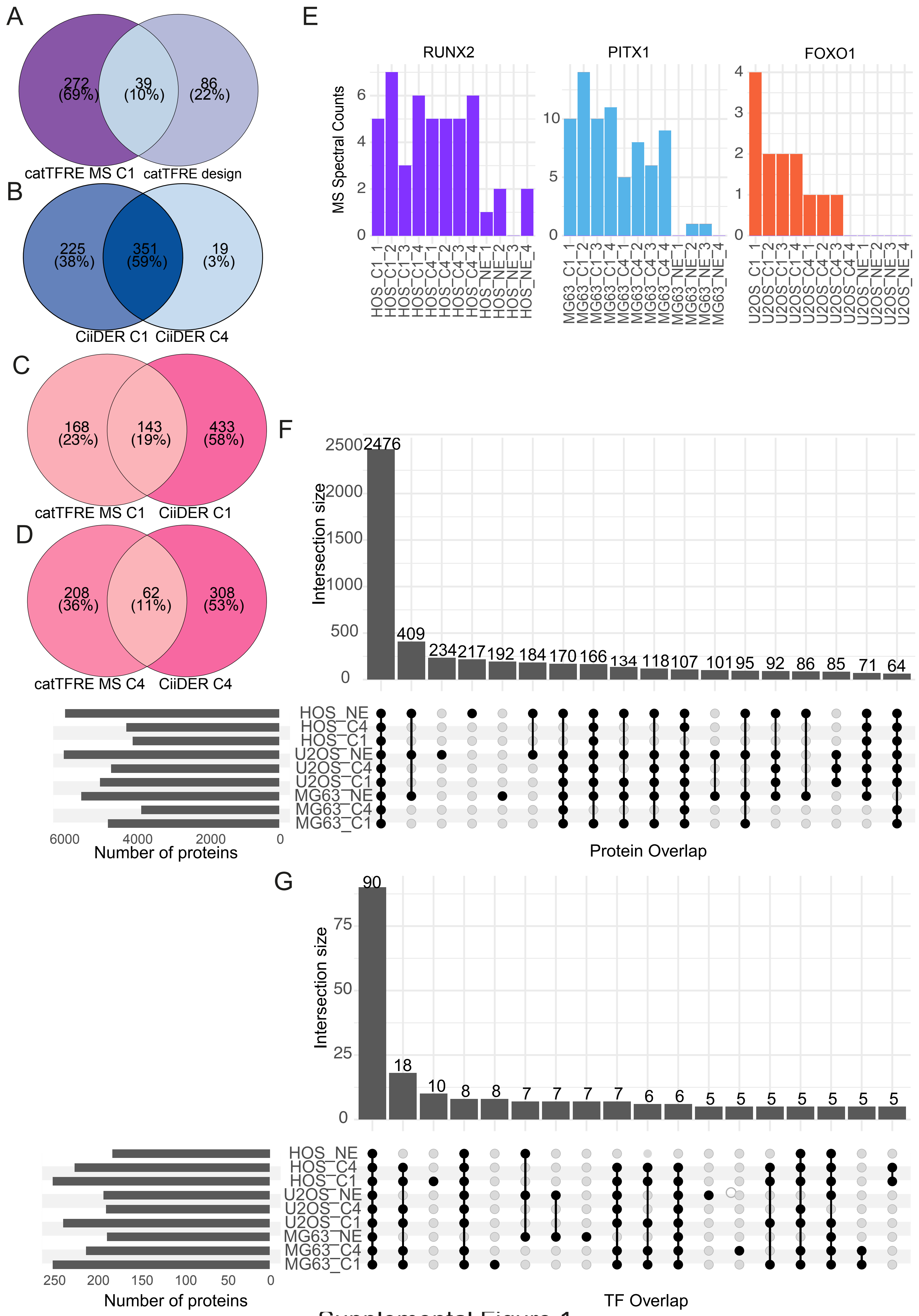


Supplemental Figure 1

Figure S1. catTFREs for enrichment of proteins and transcription factors compared to nuclear extracts.

A) Venn diagram of total TF detected in catTFRE MS C1 and catTFRE design of TF motifs.
B) Venn diagram of TF motifs detected by CiiDER in C1 and TF motifs detected by CiiDER in C4.
C) Venn diagram of total TF detected in catTFRE MS C1 and TF motifs detected by CiiDER in C1.
D) Venn diagram of total TF detected in catTFRE MS C4 TF motifs detected by CiiDER in C4.
E) Enrichment for TFs RUNX2 (purple), PITX (blue), and FOXO1 (orange) MS raw spectral count from NEs (NE) and catTFREs (DNA sequences C1 and C4) from MG63, HOS and U2OS.
E) and F) Overlapping proteins detected in NE from different OS cell lines and pulled down by catTFREs (DNA sequences C1 and C4). Results combined from four biological replicates of all proteins (in B) and TFs (in C). Source data are presented in Table S1.

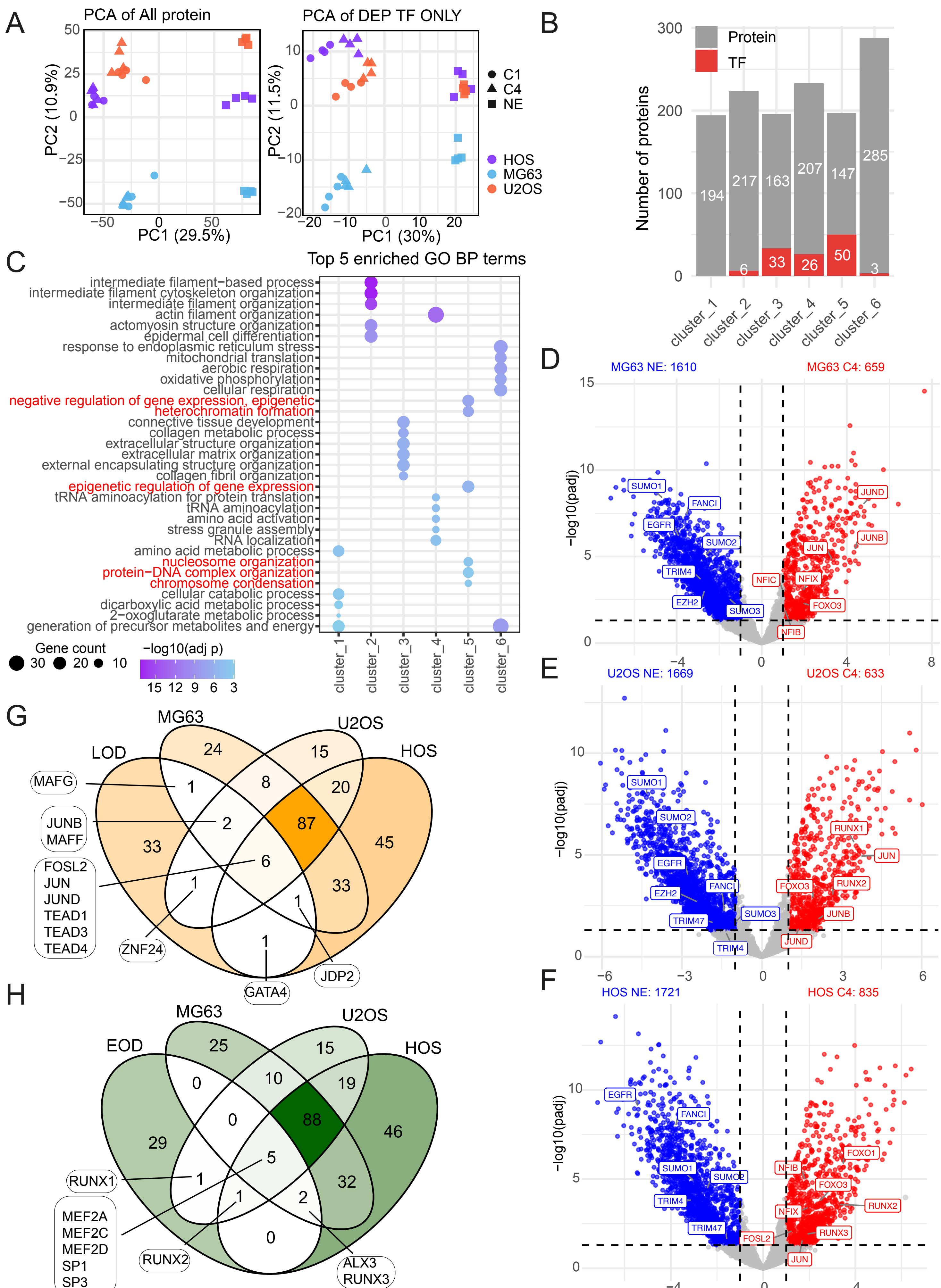


Supplemental Figure 2

Figure S2. Differential protein analysis.

A) Principal component analysis of the top 500 proteins (left) and top 200 TFs (right) in catTFREs spectral counts. Colours representing cell lines HOS (purple), U2OS (red) and MG63 (blue) and shapes representing C1 (circle), C4 (triangle), and NE (square).

B) Number of proteins and TFs in each cluster of differential protein analysis in Figure 2A, red colour representing TFs and grey colour representing other proteins.

C) Top 5 of gene ontology analysis of biological processes for proteins in clusters 1-6 in Figure 2A. Cluster 5 represents the catTFREs proteins that were highlighted in red.

D, E, and F) Volcano plots of differential protein expression (DEP) analysis between C4 and NE in MG63 (in D), U2OS (in E), and HOS (in F), abs(log2FoldChange) > 1 and a p-value < 0.05. Significant proteins detected are coloured in blue (NE) and red (C4) with a selection of key proteins highlighted.

G) A Venn diagram of all TFs significantly enriched in catTFREs C1 and C4 over NE, for MG63, U2OS, and HOS. These MS detected TFs are overlapped with a list of TF motifs correlating with late These MS detected TFs are overlapped with a list of 45 TF motifs correlating with late osteoblast-derived state (LOD) derived from López-Fuentes et al., 2025 [28].

H) A Venn diagram of all TFs significantly enriched in catTFREs C1 and C4 over NE, for MG63, U2OS, and HOS. These MS detected TFs are overlapped with a list of 39 TF motifs correlating with early osteoblast-derived state (EOD) derived from López-Fuentes et al., 2025 [28].

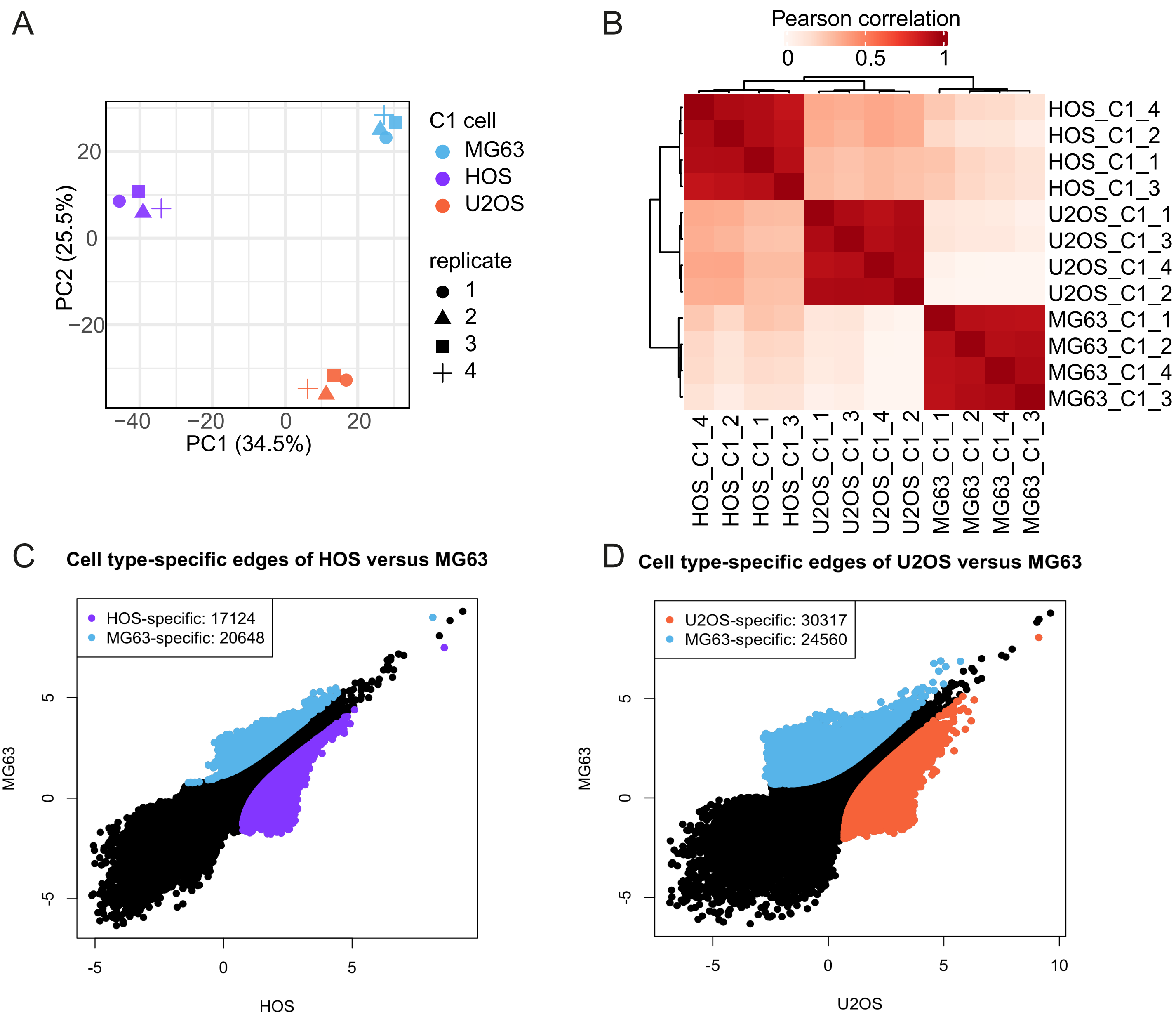


Supplemental Figure 3

Figure S3. catTFREs C1 MS comparative analysis between cell lines.

A) Principal component analysis of the top 500 all proteins from the catTFREs construct C1 of osteosarcoma cell lines. OS cell lines are represented by different colours, HOS (purple), MG63 (blue), and U2OS (orange). The shapes (circle, triangle, square and cross) are used to visualize the biological replicates.

B) Pearson correlation of raw count data from the catTFREs enrichment of C1. The heatmap represents lower correlation in white and higher correlation in red (catTFREs).

C) Cell-type-specific edges of HOS/MG63 comparison, illustrating by colour that represents the cell lines (blue, MG63 and purple, HOS). The edge differences (calculated by z-core) were predicted, weighted, and filtered from edge pools with a confidence level of at least 75%chance of being both real and different.

D) Cell-type-specific edges of U2OS/MG63 comparison, illustrating by colour that represents the cell lines (blue, MG63 and orange, U2OS). The edge differences (calculated by z-core) were predicted, weighted, and filtered from edge pools with a confidence level of at least 75% chance of being both real and different.

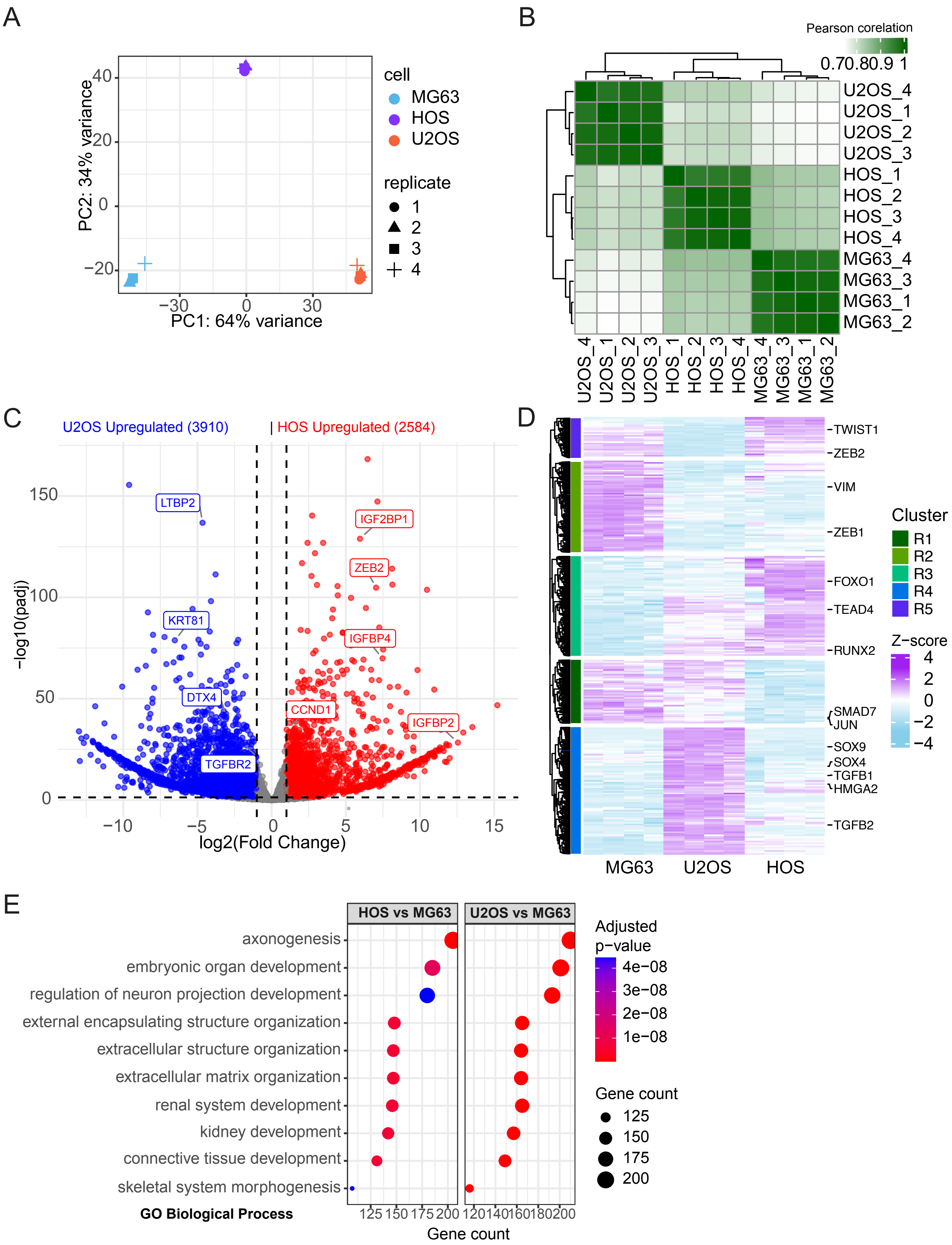


Supplemental Figure 4

Figure S4. Integrating RNA-seq and catTFREs MS to identify transcription factor candidates that regulate higher malignancy OS.

A) Principal component analysis of the four osteosarcoma cell lines RNA-seq replicates. OS cell line and biological replicates are represented by different colours and shapes, respectively.

B) Pearson correlation of the raw count of RNA-seq data from osteosarcoma cell lines. The heatmap represents lower correlation in white and higher correlation in green of the RNA-seq datasets.

C) Differential gene expression (DEGs) in the comparison of HOS versus U2OS. Statistically significant higher expression of HOS and U2OS genes are highlighted in red and blue, respectively.

D) Unsupervised clustering of DEGs from raw counts RNA-seq data set. Five clusters were identified (R1-R5), and TFs are represented for each cluster (on the right).

E) Top 10 overlapping GO terms from clusterProfiler analysis using differentially expressed gene data in Fig. 4A (U2OS/MG63 RNA-seq) and Fig. 4B (HOS/MG63 RNA-seq).

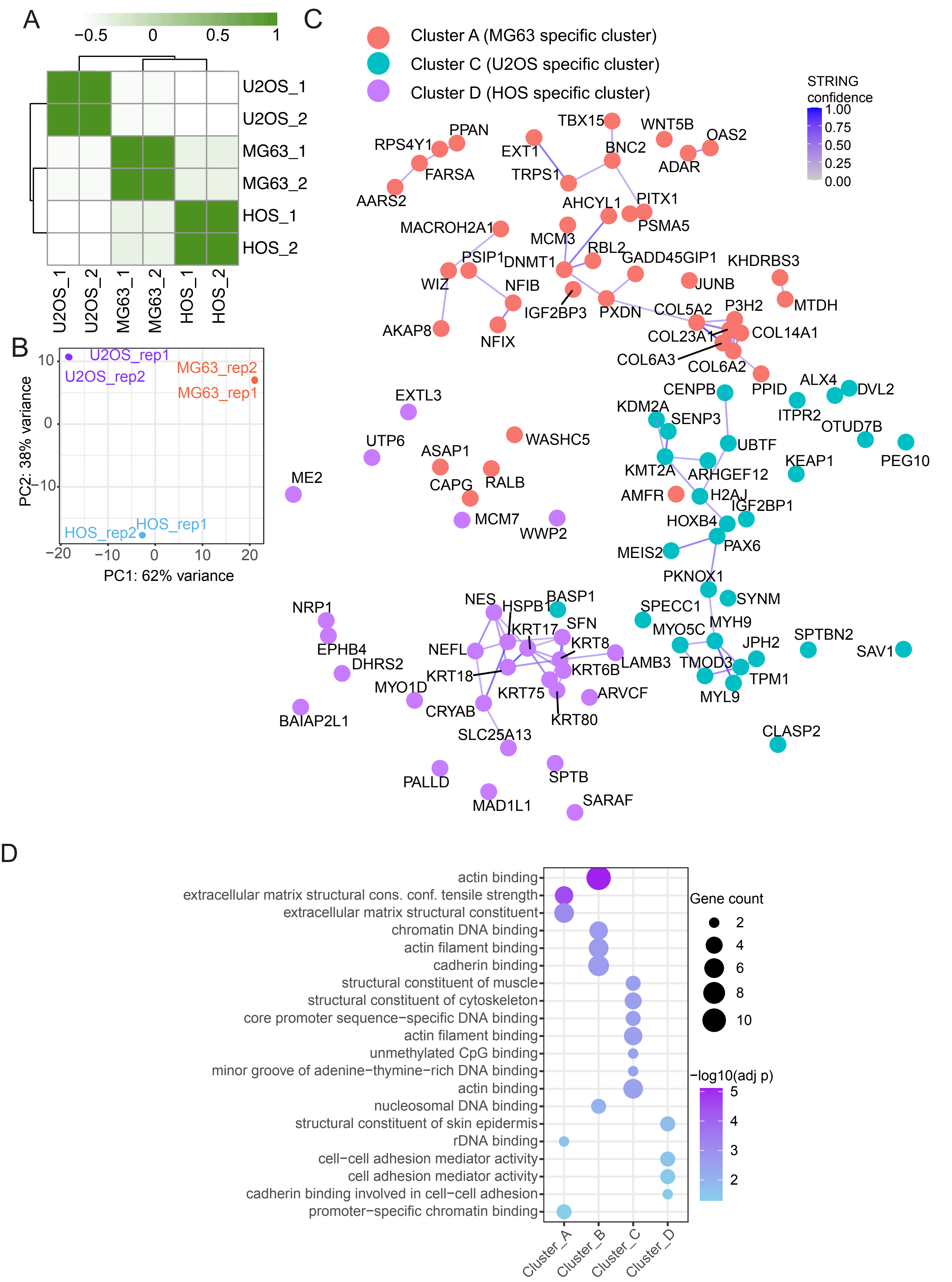


Supplement Figure 5

Figure S5. Overlapping catTFREs MS and transcription factor motif binding in chromatin accessibility.
A) An overview of ATAC-seq data of OS cell lines with Pearson correlation
B) Principal component analysis of ATAC-seq data of OS cell lines.
C) Protein–protein interactions of MG63 (red), U2OS (purple) and HOS (turquoise) specific clusters from Fig. 5A predicted using the StringDb database. Each cluster represents the protein interaction network individually, with a STRING confidence of at least 0.4.
D) Gene ontology (GO) analysis of protein clusters from Fig. 5A.
